# Protein structure alignment significance is often exaggerated

**DOI:** 10.1101/2025.07.17.665375

**Authors:** Robert C. Edgar, Harutyun Sahakyan

**Author notes:** Contributing authors.

## Abstract

Machine learning has generated millions of high-quality predicted protein structures, creating a need for computationally efficient structure search algorithms and robust estimates of statistical significance at this scale. We show that unrelated proteins have a universal tendency towards convergent evolution of secondary and tertiary motifs, causing an excess of high-scoring false positive alignments. We investigate popular structure search and alignment algorithms, finding that previous methods routinely overestimate significance by up to six orders of magnitude. To address these issues, and to accommodate recent innovations in search algorithm design, we describe a novel method for estimating statistical significance. We show that its *E*-values are accurate, scale successfully with database size, and are robust against the (generally unknown) diversity of folds in the database.

We implement our approach in an online structure search service based on Reseek at https://reseek.online.

## 1 Introduction

The explosive growth of biological sequence data has revolutionized biology, transforming our understanding of life at the molecular level. Searching novel proteins against comprehensive public databases has emerged as an indispensable, foundational strategy for biological research. Recent machine-learning (ML) breakthroughs including AlphaFold (AF) (Jumper et al. 2021) and ESMFold (Lin et al. 2023) have boosted the value of low-cost sequencing by adding high-quality predicted structures of hundreds of millions of proteins (Varadi et al. 2022), enabling richer characterization and identification of more distant homologs. The canonical first step in exploiting these resources is to construct and rank alignments for a protein of interest against proteins in a database. For sequence search, the gold standard for ranking hits is by *E*-value, a measure of significance pioneered by BLAST (Altschul et al. 1997) and subsequently adopted by other search algorithms such as hidden Markov models (Karplus et al. 2005). Prior to the ML folding era, popular structure alignment methods included DALI (Holm and Sander 1998) and TM-align (Zhang and Skolnick 2005). These algorithms take roughly one to ten seconds per alignment on current commodity computers and are therefore too slow to be practical for routine searches at the scale of the AlphaFold Structure Database (AFDB, currently 214 million proteins). More recently, Foldseek (Van Kempen et al. 2024) and Reseek (Edgar 2024) achieved orders of magnitude faster structure search. Foldseek and Reseek represent structures by sequences in a 20-letter alphabet (3Di) and Mega, an alphabet with ~10^11^ discrete states, respectively. Here, we describe a new framework for calculating principled measures of significance for structure alignments, which is designed to accommodate recent innovations in search algorithms and enable scaling to the ML folding era.

### 1.1 Distinguishing TPs from FPs

Alignment scores are designed to sort true positives (TPs) ahead of false positives (FP). However, in practice this sorting is far from perfect and setting an appropriate score threshold is challenging, as illustrated by Fig. 1 which shows TP and FP distributions of TM scores. A large majority of alignments are FPs, while most TPs have low scores which overlap the bulk of the FP distribution. At TM=0.6, the number of TPs and FPs is approximately equal. Setting a cutoff at TM=0.6 gives FP rate of 25% and false-negative (FN) rate of 75%. Increasing the cutoff reduces the number of FPs at the expense of further increasing the high FN rate. Conversely, decreasing the cutoff rapidly inflates the already high FP rate with only a gradual improvement in TPs. Similar distributions are seen with other scores (Supp. Fig. S1) Thus, the desirable goals of detecting TPs and suppressing FPs are in strong conflict, motivating the use of *E*-values to set a threshold on the expected number of FPs.

**Fig. 1:**
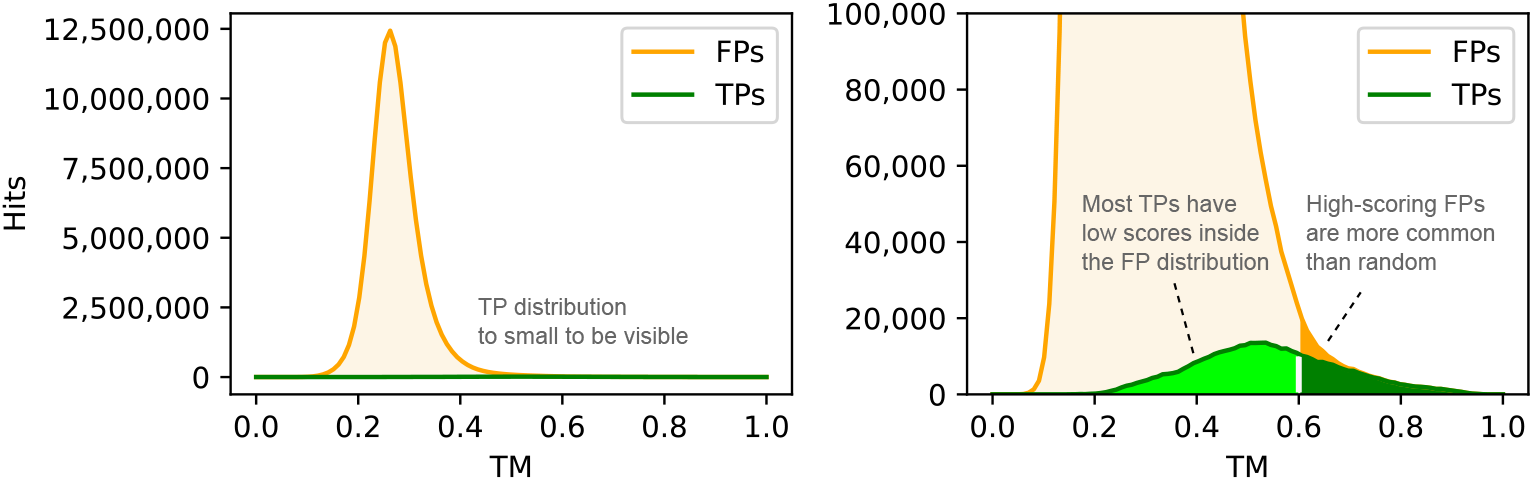
Distribution of TM-align scores Scores of TPs and FPs from TM-align all-vs-all search of SCOP40 using superfamily as a truth standard, shown as a stacked histogram binned into TM score intervals of size 0.01. Displaying the full *y* axis (left) shows that the large majority of alignments are FPs. At TM=0.6 the numbers of TPs and FPs are approximately equal. Truncating the *y* axis so that TPs are visible (right) shows that most TPs have low scores. See Supp. Fig. S1 for corresponding plots of DALI Z score and Foldseek E-value.

### 1.2 Statistical significance

The textbook approach to estimating significance (Diez et al. 2012) requires a score for each observation and a so-called null model which characterizes scores assuming that the tested relationship is absent. In practice, a null model is implemented by specifying a procedure for generating scores (here, alignment scores) at random. The significance of an alignment with score ≥ *s* is then quantified as the probability that an alignment with score ≥ *s* would be generated by the chosen null model; this probability is called the *p*-value of *s*. Sometimes, observations with *p <* 0.05 are considered significant, though this convention is arbitrary and notoriously controversial (e.g., (Amrhein et al. 2019)). For sequence search, a commonly-used null model aligns random sequences (Karlin and Altschul 1990). Then, the *p*-value of a hit with score ≥ *s* is the probability that a random sequence with the same length as the query would have a hit with score ≥ *s* to a random sequence having the same length as the search database.

### 1.3 E-values

A severe multiple test problem (Bender and Lange 2001) arises from searching a large database—a single observation with low *p*-value is unlikely by chance, but we may nevertheless see low *p*-values by chance if we repeat a given test multiple times. For example, if we find hundreds of alignments each with *p* = 1*/*100, then we should expect many of them to be false positives (FPs). This concern can be addressed by calculating an *E*-value for the alignment score *s*, i.e. an estimate of the number of FP errors per query (FPEPQ) that would be obtained by setting a score threshold of *s*. It is calculated by determining the average number of alignments with score ≥ *s* generated by the null model with a database of the same size. If *E* ≪ 1, then most null model searches have no hits with score ≥ *s*, and this justifies considering *s* to be significant. If a query is aligned to *D* proteins in a database and a given alignment score ≥ *s* has *p*-value *p*(*s*), then the *E*-value of this alignment is the probability of observing a score ≥ *s* in one observation multiplied by the number of observations, i.e.

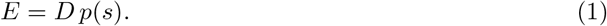

### 1.4 Gumbel distribution

The probability density function (PDF) of a Gumbel (Gumbel 1935) Extreme Value Distribution distribution is

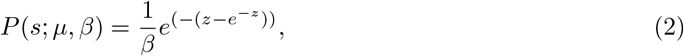

where *β* is scale, *µ* is location, and *z* = (*s* − *µ*)*/β*. Karlin-Altschul (K-A) statistics show (Karlin and Altschul 1990) that scores for gapless Smith-Waterman (Smith et al. 1981) (S-W) alignments between random sequences follow a Gumbel distribution. Empirically, scores for gapped S-W alignments are also well-approximated by a Gumbel distribution (Altschul et al. 1997).

### 1.5 Previous significance measures for structure alignments

Significance estimates from previous structure alignment algorithms are obtained by fits to Guassian or Gumbel distributions. CE (Shindyalov and Bourne 1998) and DALI report Z scores obtained from the mean 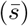 and standard deviation (*σ*) for a Gaussian fit, then

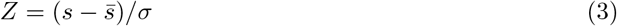

is the distance of *s* from the mean expressed as a number of standard deviations. FATCAT (Ye and Godzik 2004) fits scores to a Gumbel distribution and calculates a *p*-value similarly to K-A statistics; a similar approach for the TM score is described by (Xu and Zhang 2010). Foldseek models scores by Gumbel distributions where *β* and *µ* are predicted separately for each query using a neural network. For an alignment with score *s*, a *p*-value *p*_foldseek_(*s*) is estimated from the query’s Gumbel distribution, then the *E*-value for a database of size *D* is calculated using a modified version of Eq. 1,

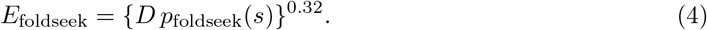

Foldseek also reports an estimated probability that an alignment with the same score would be a TP in an all-vs-all search of its training database (`prob` field).

### 1.6 Filters

In contrast to previous methods, which unconditionally align every pair of structures, Foldseek and Reseek use *filters*, heuristics that optimize speed by reducing the number of alignments (Fig. 2). We distinguish between *intrinsic filters*, which use only information about a pair of structures to decide whether the pair should be discarded, and *data-dependent filters*, where the outcome depends on other factors such as the size and composition of the search database. Both Foldseek and Reseek employ filters which use short, fixed-length words (*k*-mers in the 3Di and *µ* structure alphabets, respectively; here *µ* is unrelated to the location parameter of a Gumbel) to find diagonals with score above a threshold (*d*). Database structures (*targets*) passing the diagonal test are stored in a buffer of fixed length *K*, where *K* is set by the `max-seqs` option of Foldseek or the `rsb size` option of Reseek. If there are *> K* targets with diagonal score ≥ *d* for a given query, then the buffer overflows (by design), and only the top *K* targets (by *d*) are aligned to that query. The rate of overflows depends on the database size—if there are more targets, the buffer is more likely to overflow. For very large databases, most targets with score ≥ *d* are discarded due to buffer overflow and the resulting speedup is substantial. This type of diagonal filter is therefore data-dependent, having an increasing impact on the alignment score distribution as databases grow larger. Reseek employs an additional filter (mu) which discards targets if their alignment in the *µ* alphabet is below a threshold. The mu filter gives a useful speedup because *µ* is much smaller than the Mega alphabet used for the final alignment, enabling a fast implementation based on SIMD instructions. Mu is an intrinsic filter because it is applied unconditionally to every query-target pair which was passed by the diagonal filter. Mu is strongly biased to discard pairs with low alignment scores, thereby biasing reported alignments towards higher scores. Filters tend to reduce FPEPQ by reducing the number of opportunities for false positives. Consider for example a filter which discards 99% of targets at random. This filter would be undesirable in practice, but regardless the expected number of false positives will be reduced by a factor of 100 and the *E*-value should therefore be reduced by the same factor. A robust *E*-value estimate must therefore consider the effects of filters.

**Fig. 2:**
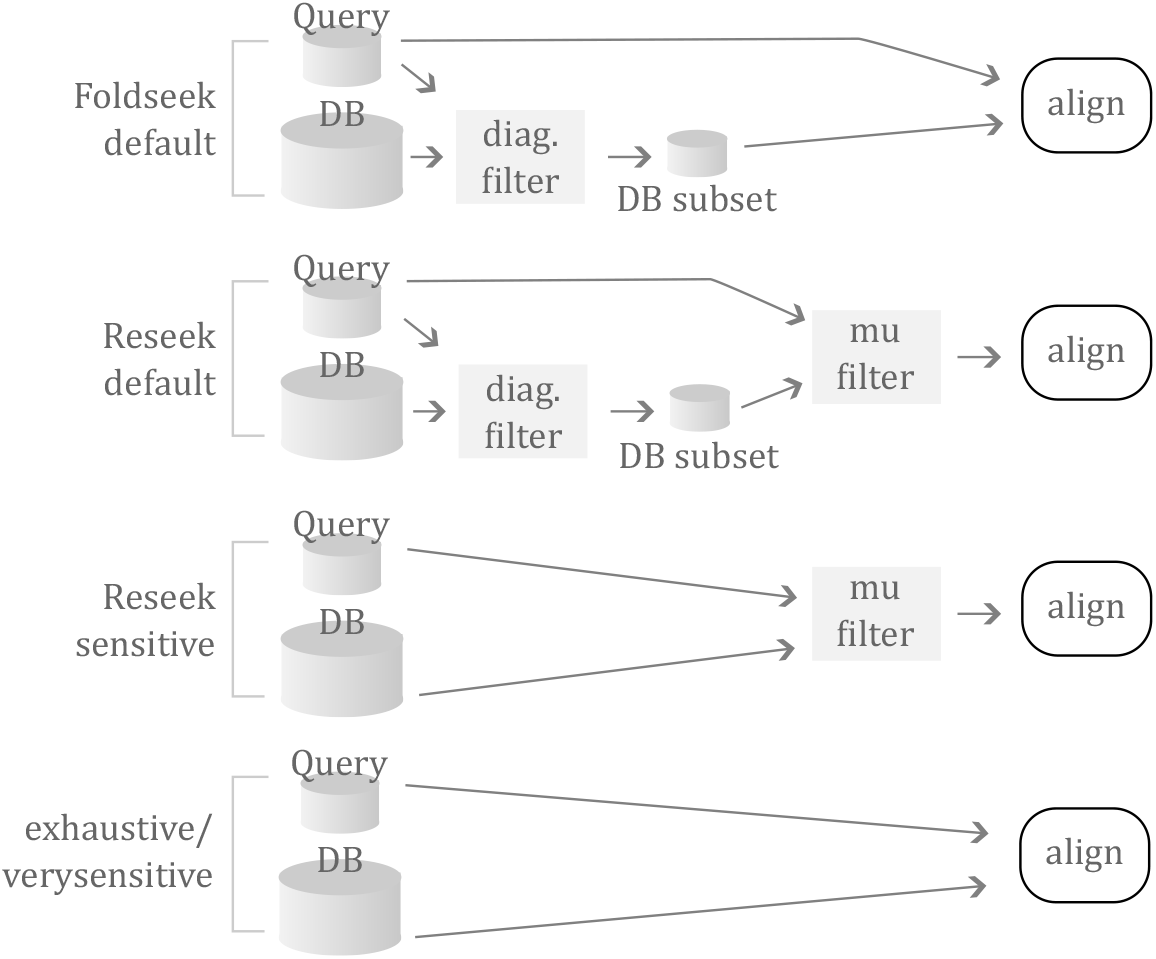
Workflows of the Foldseek and Reseek algorithms. Foldseek with default options and Reseek with the `fast` option use a diagonal filter (diag.) which checks for two *k*-mers on the same diagonal and stores the top *K* targets for each query structure. By design, this buffer tends to overflow with large databases, reducing the number of alignments. Reseek with `fast or sensitive` also applies a filter (mu) which performs a fast, approximate alignment, discarding hits below a threshold; this filter is applied unconditionally to any query, target pair. With the `verysensitive` option of Reseek or `exhaustive-search` option of Foldseek, filters are disabled so that all queries are aligned to all targets.

### 1.7 Structure search null model

Conceptually, the null model proposed here (Ø) simulates the workflow of a structure search pipeline (Fig. 3). It is designed to model any search algorithm with an alignment step, allowing intrinsic and/or data-dependent filters. The query and database are each characterized by their *alpha diversity*, i.e. by a frequency distribution over distinct structures. By our definition, an alignment is a TP if the structures are homologous, otherwise a false positive. An alternative truth standard can be accommodated in this framework providing there is a suitable trusted reference. The first step in the simulation samples a query structure *q* and target structure *t* at random and discards the pair if it is a TP. The pair is tested by the filters, if any, employed by the pipeline, and aligned if passed by those filters. Running this model repeatedly yields a distribution *P*_Ø_(*s*|FP) of alignment scores for false-positive pairs passing the filters, which is then used to calculate *p*-values and *E*-values. Implementing this abstract model in practice raises a number of challenges, including modeling alpha diversities, identifying false-positive pairs, and modeling filters. Briefly, these challenges are addressed as follows (Fig. 4). We show empirically that the score distribution of false positives is independent of alpha diversity to a good approximation, which allows us to use any trusted reference database as the initial source for sampling, giving a *universal* (i.e., reference database- and alpha diversity-independent) distribution *P*_*u*_(*s*|FP). An intrinsic filter (IF) is modeled by *P*_IF_(+|*s*), the probability that a pair with alignment score *s* is passed by the filter; this probability is independent of the search data (by definition), so can be measured once and hard-coded thereafter. A data-dependent filter (DDF) is modeled by its pass probability *P*_DDF_(+|*s*, **Q, D**) which is measured on the search database **D** using a representative query dataset **Q**. The score distribution for the complete pipeline is then

**Fig. 3:**
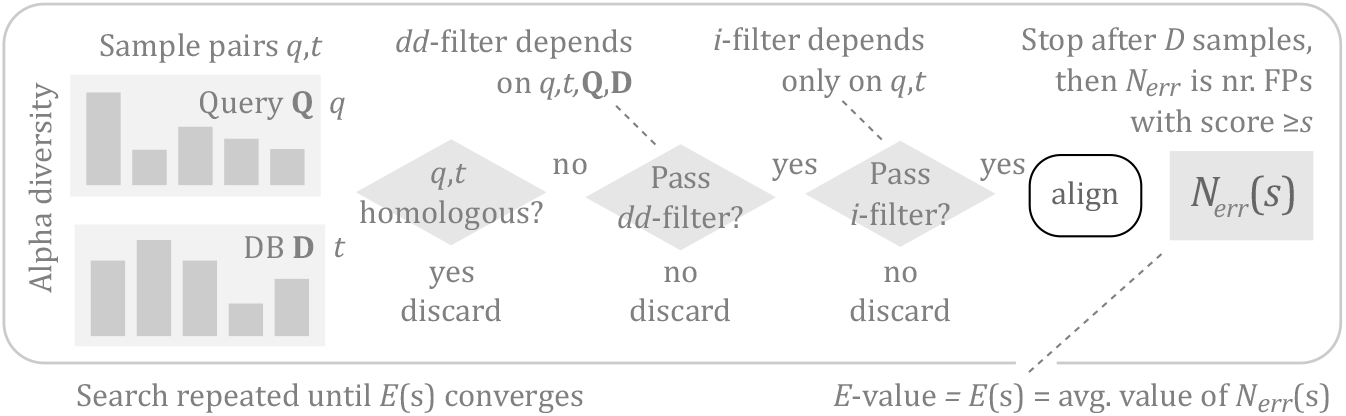
Abstract null model. In our approach, an idealized (*abstract*) null model (Ø) simulates search algorithm workflows like those seen in Fig. 2. The query (**Q**) and database (**D**, containing *D* structures) are each represented by an alpha diversity, i.e. by a frequency distribution over distinct structures. In general, these diversities may be different, modeling a scenario where, say, virus capsid proteins are searched against a database of eukaryotic proteins. Pairs of structures *q,t* are sampled at random from **Q** and **D** respectively. A pair is discarded if *q* and *t* are homologous. Pairs are tested and potentially discarded by a data-dependent (*dd*-) filter, where the outcome depends on other pairs passing through the filter, and by an intrinsic (*i*-) filter, which depends only on *q* and *t*. If a given type of filter is not used by an algorithm, the corresponding null model filter passes the pair unconditionally. If the pair passes all filters, it is aligned to obtain the score ≥ *s*. When *D* pairs have been sampled, the total number of alignments *N*_*err*_(*s*) with score *s* is calculated. This procedure generates *D* alignments, thereby modeling the search of a single query structure against the database. The search is repeated until the average value of *N*_*err*_(*s*) converges; this average is the expected number of false positives per query with score at least *s*, i.e. the *E*-value according to this null model.

**Fig. 4:**
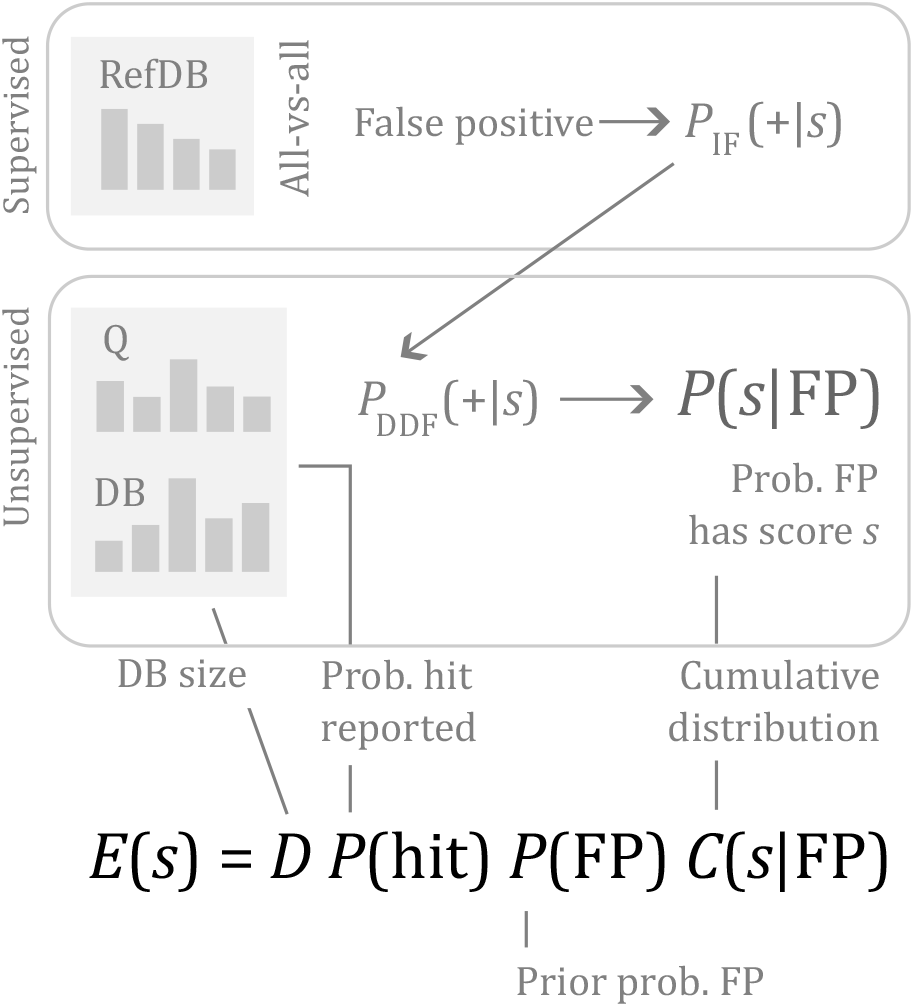
Empirical null model Implementing the abstract null model shown in Fig. 3 raises a number of practical issues. This figure sketches our solution, which factors the model into a supervised module using a gold-standard reference database (here generically called RefDB) for homology, and an unsupervised module which analyzes hits to the search database **D** used in practice (e.g., AFDB50) together with a representative query dataset **Q**. Arrows represent the conceptual flow through the model, which is implemented by multiplying probabilities *P* (step1) *P* (step2)…because a FP hit must progress through all steps to be reported. Filters are modeled by estimating the probability *P* (+|*s*) that a *q,t* pair with score *s* will be passed by the filter. Conceptually, an alignment with score *s* is discarded stochastically by a filter with probability *P* (+|*s*). For an intrinsic filter, *P*_IF_(+|*s*) depends only on the score and can therefore be measured once and hard-coded thereafter; for convenience, we choose to measure it on the reference. For a data-dependent filter, *P*_DDF_(+ |*s*, **D, Q**) is measured using **D** and **Q** for each required combination of **D** and **Q**. *P* (reported) is the probability that a randomly chosen pair from **D**,**Q** has a reported alignment; this is straightforwardly measured as the total number of reported hits divided by the number of possible hits |**Q**|×|**D**|. *P* (FP) is the prior probability that a randomlychosen pair from **D**,**Q** is a false positive. We set *P* (FP) = 1 for comprehensive search, because most randomly-selected *q,t* pairs will be FPs in typical searches. For a filtered search, we set *P* (FP) = 0.5 because in general we have no prior expectation that a randomly-chosen hit which has passed the filters is more likely to be a TP or FP.

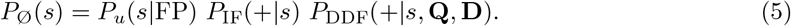

The pass probability for a filter can be measured using unsupervised training, i.e. without knowing which *q,t* pairs are false positives, and can therefore be measured on any search database. In this way, the simulation is factored into tractable modules: a supervised module that is independent of the search database size and composition, and an unsupervised module which can be implemented on any database used in practice, as shown in Fig. 4. The *p*-value according to Øis the probability that the score of a FP pair passed by the filters is ≥ *s*, which is calculated as the sum 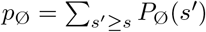. The corresponding *E*-value is *E*_Ø_(*s*) = *D p*_Ø_(*s*) per Eq.(1),

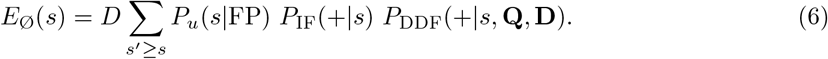

If there are no filters, then Eq.(6) becomes 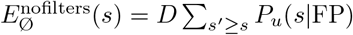,or

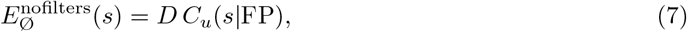

where *C*_*u*_(*s*|FP) is the cumulative distribution of *P*_*u*_(*s*|FP).

### 1.8 Gold standards for false positives

The SCOP database (Murzin et al. 1995) classifies protein domains into folds and superfamilies. Folds are somewhat arbitrary categories for the overall conformation (see Discussion). Folds are divided into superfamilies. Domains assigned to the same superfamily are believed to be homologous, while different superfamilies are ambiguous—similarities between two superfamilies in the same fold may be similar due to convergence rather than homology. SCOP40 and SCOP95 are subsets of SCOP clustered so that the maximum amino acid (a.a.) sequence identity is 40% and 95% respectively. CATH (Orengo et al. 1997) classifies domains into topologies (roughly equivalent to folds in SCOP), which are similarly divided into superfamilies. As with SCOP, domains in the same CATH superfamily are believed to be homologous, whereas different superfamilies in the same topology may be convergent. We consider superfamily assignments as our gold standard so that a pair *q,t* is a false positive if *q* and *t* are assigned to different superfamilies. This follows (Edgar 2024), while other papers have used different standards (see Supplementary Material section “Truth standards” for discussion). Unless noted otherwise, results here are reported using SCOP40 v1.75 for consistency with previous literature, especially (Van Kempen et al. 2024). We also created a subset SCOP40 which we believe is better suited for assessing homolog detection accuracy and training a null model (see Supplementary Material section “SCOP40 curation”). This subset, which we call SCOP40c, was used to train defaults in Reseek v2.6.

## 2 Results

### 2.1 FP score distributions

Fig. 5 shows a Gumbel fit to Foldseek and Reseek S-W scores together with TM scores for TM-align, measured on SCOP40 FPs, showing a good fit to the bulk of the distribution. For S-W scores, a Gumbel fit is predicted by K-A statistics for random sequences. For TM scores, a good fit to Gumbel was observed for the bulk of the distribution in (Xu and Zhang 2010). However, when viewed on a logarithmic scale, a large excess of observed compared to fitted scores is apparent in the high-scoring tail. If a null model assumes a Gumbel distribution, this excess will cause significance of high-scoring alignments to be overestimated.

**Fig. 5:**
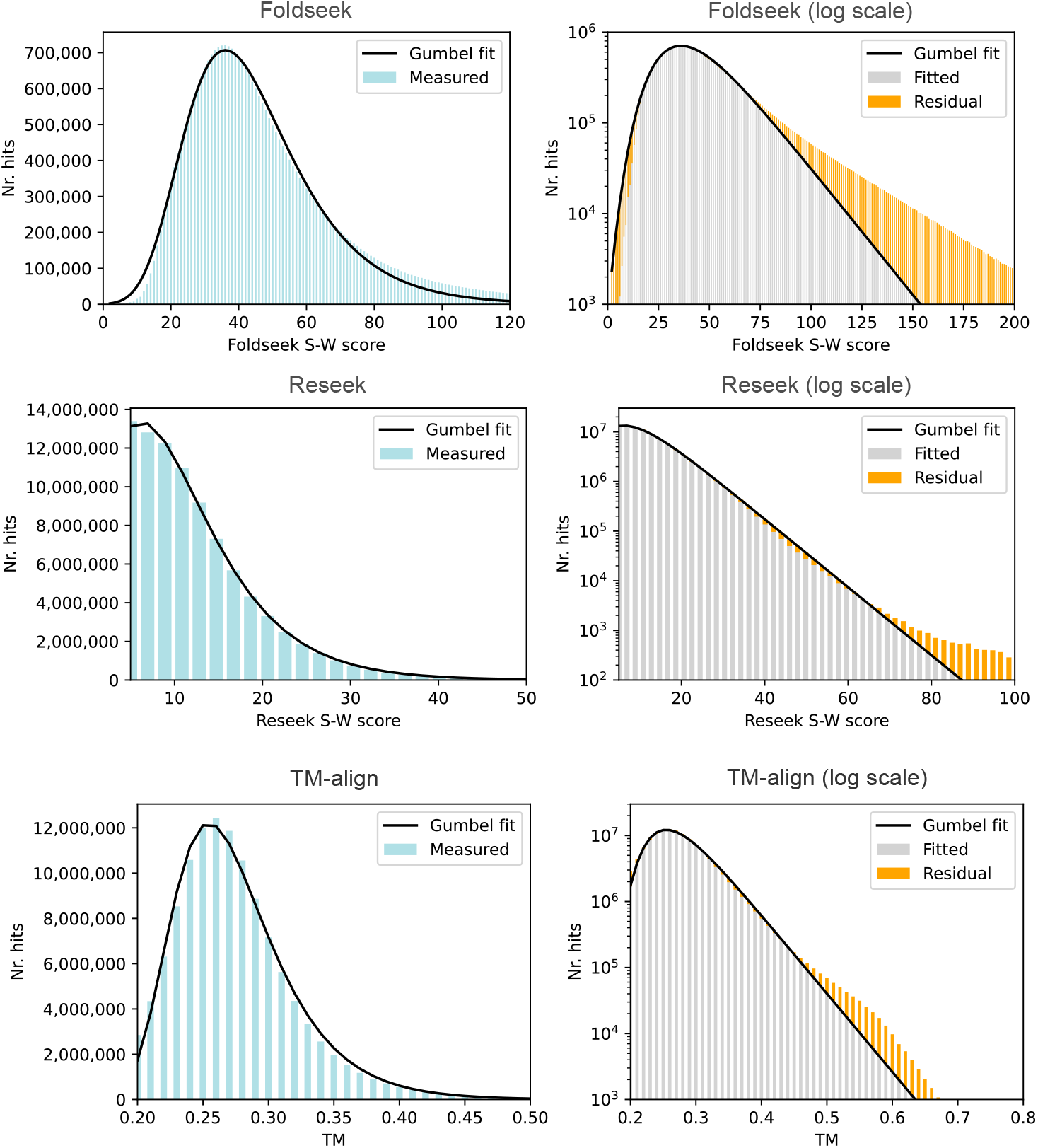
Fit to Gumbel distribution Gumbel fits to Smith-Waterman and TM scores of FPs according to the SCOP40 reference, using linear (left) and logarithmic (right) scales for the *y* axis.

### 2.2 Entropy of 3Di sequences

Repetitive (*low-complexity* or *low-entropy*) regions have runs of one letter (e.g. `VVVV`…) or a few letters (e.g. `QNQQQNQQNNN`…). Such regions may align between unrelated proteins, giving higher scores than would be expected for random sequences. We found that 1,084,741 / 1,949,312 (55.6%) 3Di letters in SCOP40 are masked by the segmasker algorithm for finding low-entropy regions (Wootton and Federhen 1993). Fig 6 shows Gumbel fits to S-W alignments in the 3Di alphabet. In the top panels, all domains are aligned to all reversed sequences. There is an excess of high-scoring alignments which is largely eliminated by deleting segmasked regions (bottom panels), confirming that low-complexity 3Di sequences cause an excess of high-scoring alignments compared to a Gumbel fit. See also Fig. 7. Similarity of low-complexity sequences in unrelated sequences is due to convergent evolution, by definition.

**Fig. 6:**
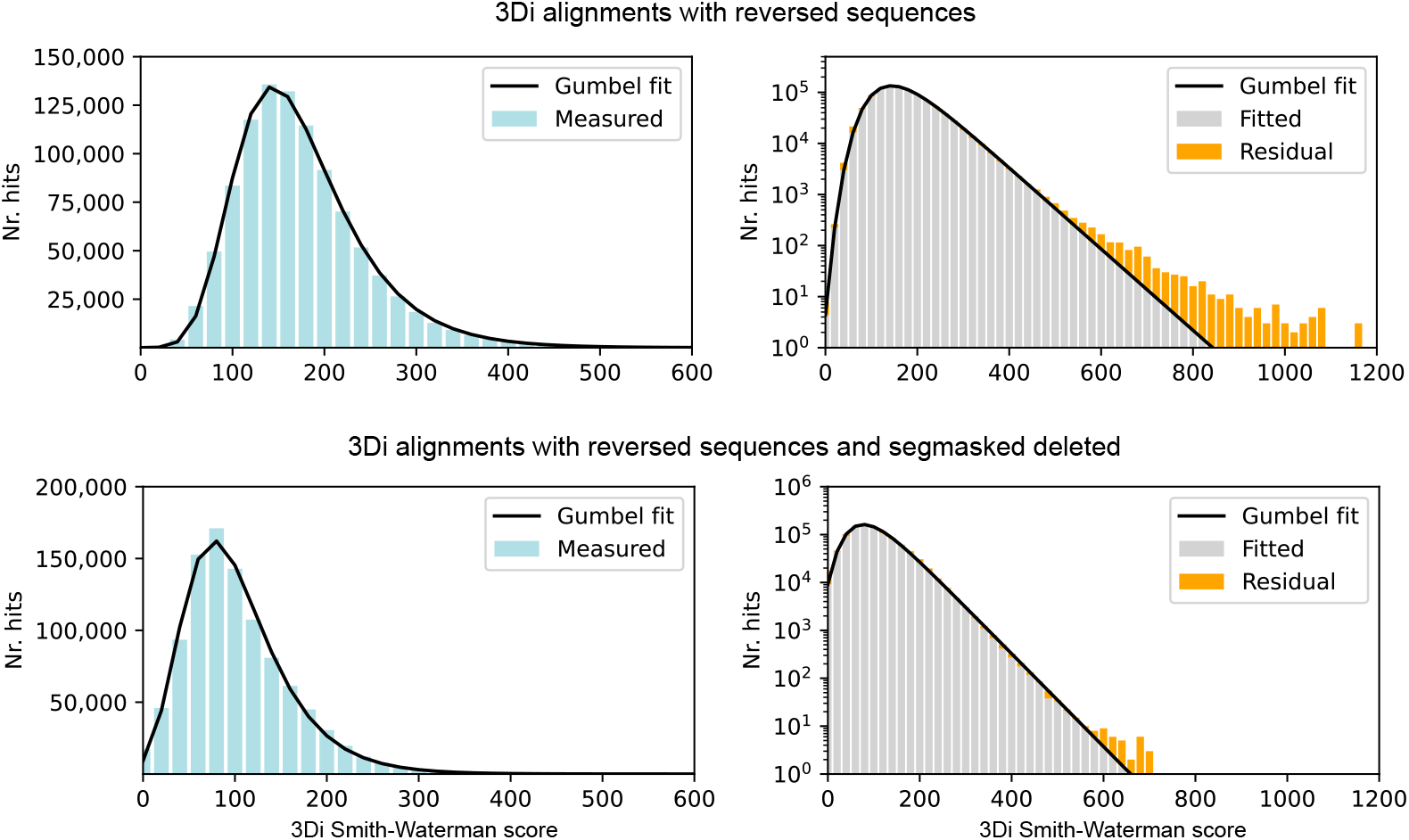
Fit to Gumbel distribution for 3Di alignments Fits to a Gumbel distribution for Smith-Waterman scores of all SCOP40 3Di sequences against all reversed SCOP40 3Di sequences (top) and for all SCOP40 3Di sequences with repetitive regions deleted (bottom).

**Fig. 7:**
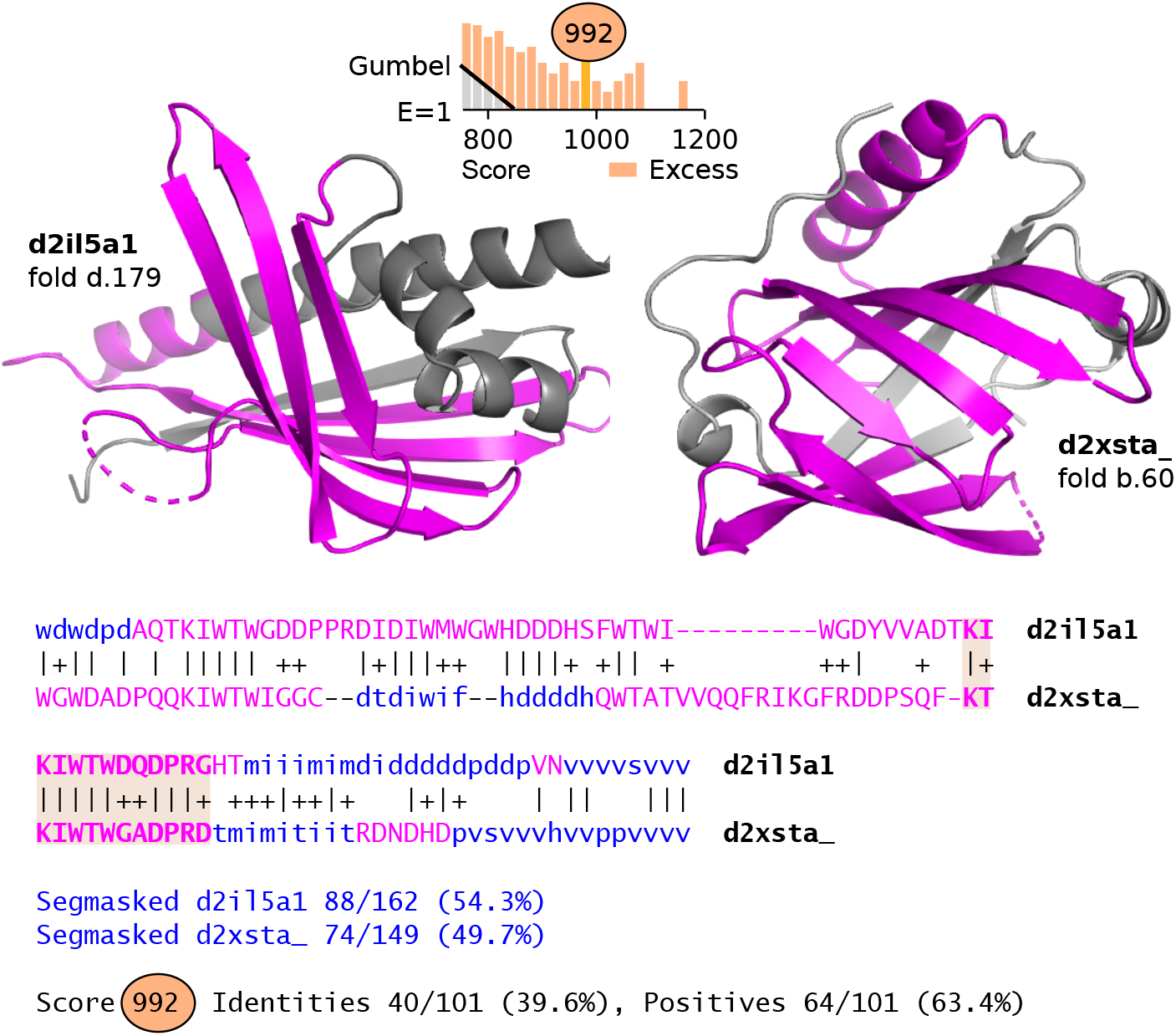
Non-random similarity between unrelated proteins The figure shows cartoon representations of unrelated domains `d2il5a1` (fold `d.179`) and d2xsta (fold `b.60`) above a Smith-Waterman alignment of their 3Di sequences.

### 2.3 Foldseek raw score distribution compared to Gumbel

The Foldseek raw score is calculated by subtracting the S-W score for the reversed query sequence from the S-W score for the original query. Foldseek fits this raw score to a Gumbel distribution to estimate its native *E*-value. However, the raw score distribution of FPs demonstrates (Fig. 8) that there is a much larger deviation from the Gumbel fit compared to S-W, especially in the high-scoring tail, which is most relevant to significance estimates. This result suggests that Foldseek E-values are likely to be substantially underestimated (confirmed below).

**Fig. 8:**
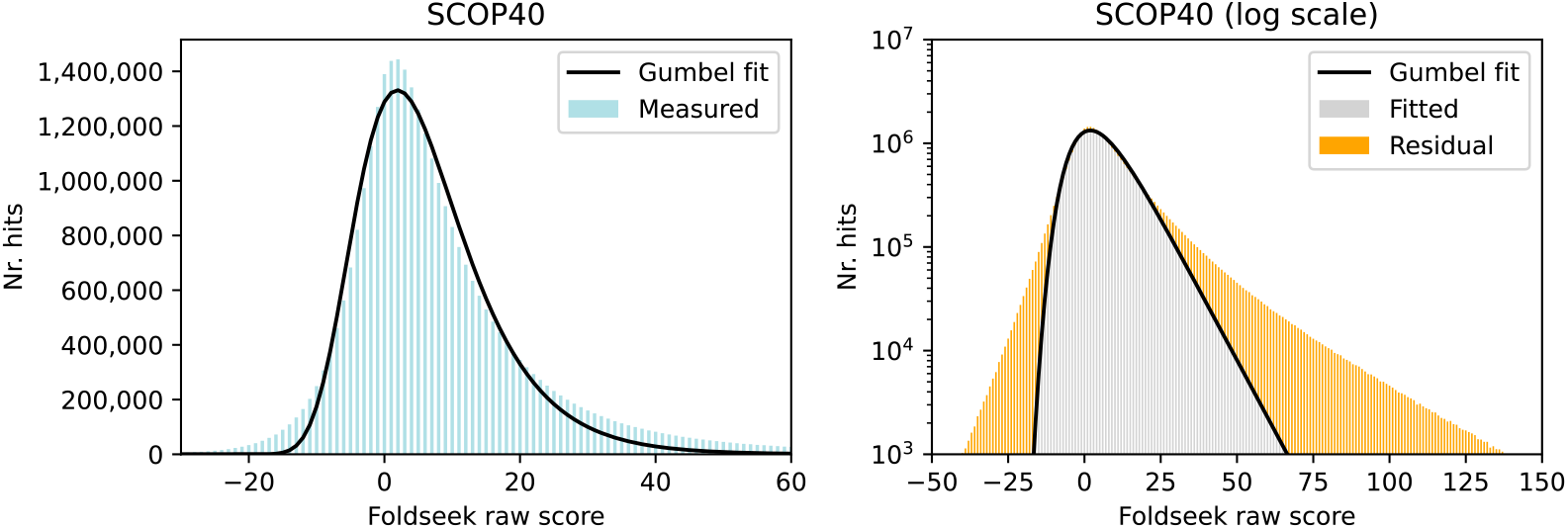
Fit to Gumbel distribution for Foldseek raw score Fits to a Gumbel distributions for Foldseek raw scores of FPs according to the SCOP40 (top) and SCOP40c references (bottom) references, using linear (left) and logarithmic (right) scales for the *y* axis. The raw score is obtained by subtracting the Smith-Waterman score for the reversed query sequence from the Smith-Waterman score for the original query. On a linear scale, the fit appears superficially reasonable, but with a logarithmic scale a large excess of observed compared to fitted is visible in both tails, especially the high-scoring tail where alignments with higher significance are found. Therefore, we expect the Gumbel fit to raw scores used by Foldseek to substantially overestimate significance of high-scoring alignments.

### 2.4 Universality of the FP score distribution

Fig. 9 shows *P* (*s*|FP) on a variety of reference databases for DALI, TM-align, and Reseek. These distributions were measured with different standards (SCOP40, SCOP40c and CATH40) and a range of *sampled* references obtained by sub-sampling and by concatenating multiple copies of the reference (super-sampling), respectively. Sub-sampling was performed either by sub-sampling domains at random (denoted SCOP40*/n* for sampling 1*/n* domains), which reduces size while leaving alpha diversity unchanged, or by sub-sampling superfamilies at random (denoted SCOP40/SF*n* for sampling 1*/n* superfamilies), which reduces size and changes alpha diversity. The variation with reference standard (SCOP or CATH), size, and alpha diversity is small, showing that *P* (*s*|FP) and *C*(*s*|FP) are universal by our definition here, i.e. independent of the database to a good approximation. Importantly, we do not see trends where the distribution changes systematically with increasing size or increasing diversity—even small systematic changes would be amplified by the leap from SCOP40 to AFDB scale (~ 20, 000×). Note that the universality of *P* (*s*|FP) does not imply the universality of *P* (FP|*s*) or *P* (TP|*s*) (see below for results showing that these distributions are not universal).

**Fig. 9:**
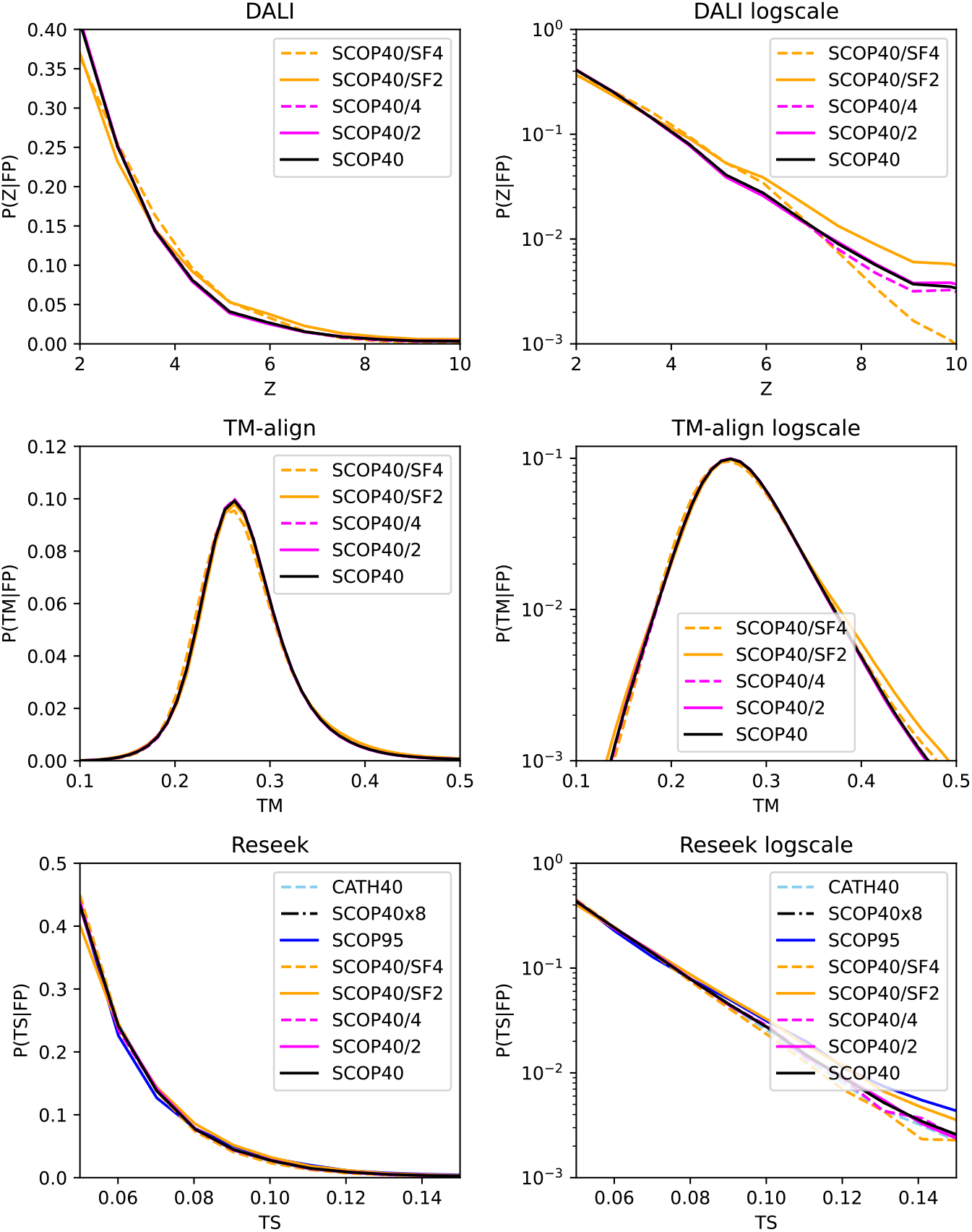
Distributions of *P* (*s*|FP) with varying references. Observed distributions of *P* (*s* FP) for DALI Z-score (top), TM score (middle) and Reseek test statistic (TS, bottom) on various references including SCOP40, SCOP95, CATH40 and sampled subsets of SCOP40. Sampling ×*n* indicates super-sampling, */n* indicates sub-sampling domains, /SF*n* indicates sub-sampling superfamilies. Linear scales are used for the *y* axis on the left, logarithmic on the right. These distributions are seen to be similar across all variations in the reference database. In particular, there is no obvious trend with increased size or increased number of superfamilies, suggesting that the distribution on a much larger database would also be similar. By contrast, the probability we would ideally like to know, i.e. *P* (TP |*s*) = 1 − *P* (FP |*s*), varies strongly with alpha diversity (Fig. 12).

### 2.5 E-values estimated by log-linear fitting

We found that the high-scoring tails of relevant probability distributions are well approximated by log-linear fits (Supplementary Fig. S2); an implementation of Øby log-linear fitting is denoted Ø_*LL*_. Methods which not use filters are modeled by Eq.(7). In a log-linear approximation using base 10, Eq.(7) becomes

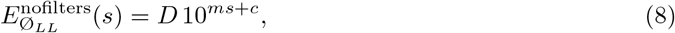

where *m* and *c* are the fitted parameters. Tables of *E*-values for DALI, TM-align and Foldseek estimated by Ø_*LL*_ are given in Supplementary Data. For example, on AFDB50 we find TM score 0.9 has *E*-value 4.4 using SCOP40 as a truth standard or 1.0 using SCOP40c. DALI Z-score 40 has *E*-value 1.2 × 10^*−*2^ (SCOP40) or 2.2 × 10^*−*7^ (SCOP40c). This last example shows that the choice of SCOP40 or SCOP40c as the truth standard can have a substantial impact on the *E*-value. Reseek implements two filters: a diagonal filter and the mu filter. The *p*-values and *E*-values reported by Reseek are based on log-linear approximations to Eqs.(5) and (6) using these fitted parameters with SCOP40c as a reference.

### 2.6 E-value accuracy and scaling

BLASTP *E*-values by Ø_*LL*_ correlate well with its native *E*-values and give a similar coverage-vs-error (CVE) plot (Supplementary Fig. S2). An *E*-value *E*(*s*) is an estimate of the number of false-positive errors per query FPEPQ(*s*) at score cutoff *s*, and its accuracy can therefore be assessed by plotting *E*(*s*) vs. FPEPQ(*s*) and comparing with an ideal *E*=FPEPQ. Results for the *E*-value estimates according to our log-linear fits are compared to measured FPEPQ for DALI, TM-align and Reseek in Supplementary Fig. S3, using SCOP40 and SCOP40c as truth standards. These results show that Ø_*LL*_ is in good agreement with measured FPEPQ, accounting well for varying database size and diversity. Plots of *E*(*s*) vs. FPEPQ(*s*) for Foldseek and Reseek are shown in Fig. 10 for a range of database sizes spanning a factor of 32 from smallest to largest, obtained by sub- and super-sampling. While Reseek *E*-values remain close to ideal across this range (Fig. 10(c)), Foldseek *E*-values are increasingly underestimated as the database size increases (Fig. 10(a)). By trial and error, we found that the following correction gave an *E*-value close to ideal,

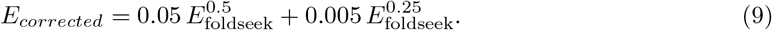

As seen in Fig. 10(b), this corrected *E*-value is comparable to Reseek’s (Fig. 10(c)) in approaching the ideal. This formula enabled us to estimate *E*_foldseek_ vs. FPEPQ(*s*) for larger databases, with results included in Table 1.

**Fig. 10:**
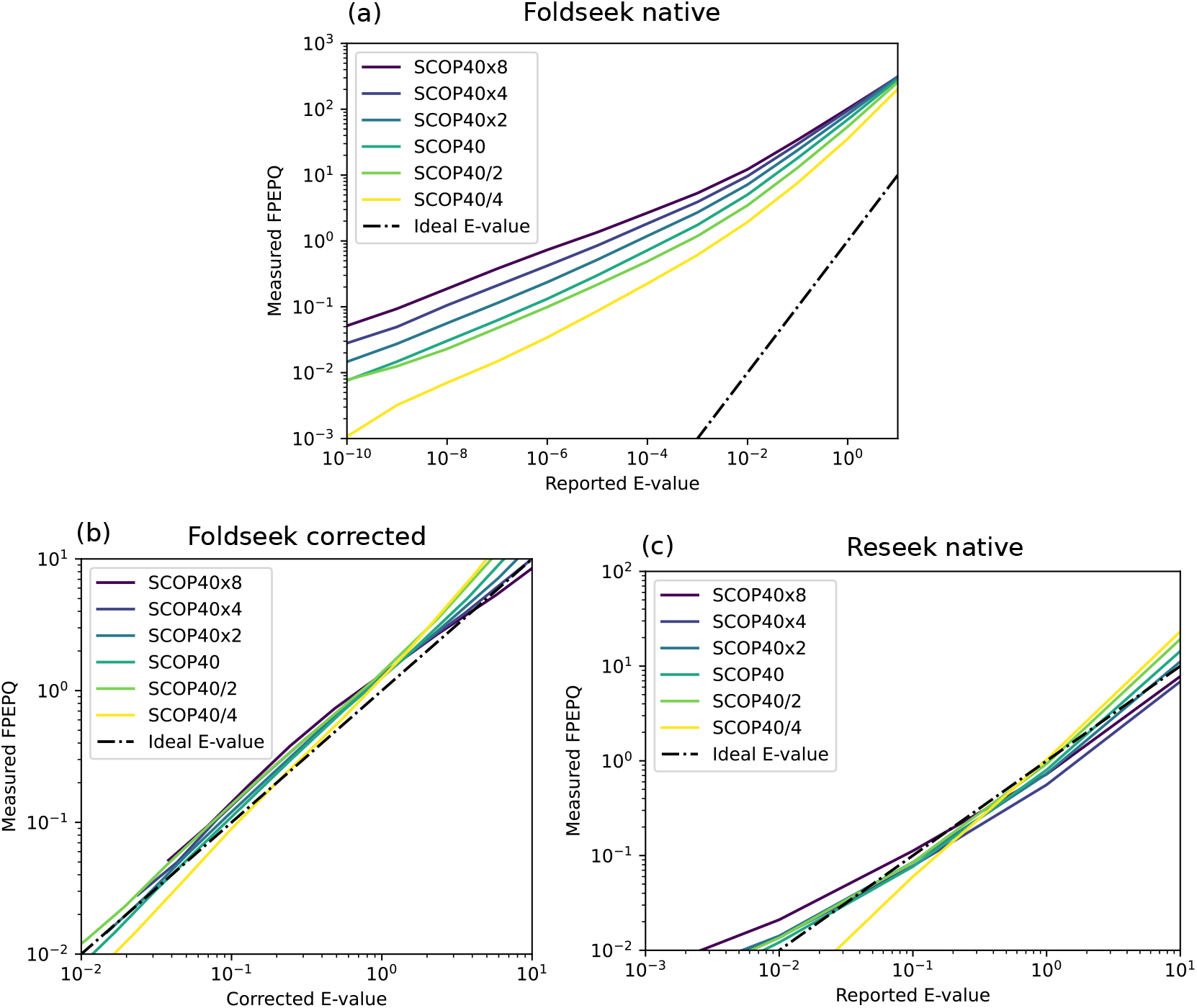
Scaling of *E*-value with database size (a) Native Foldseek *E*-value on the *x* axis against measured FPEPQ on the *y* axis. An ideal *E*-value exactly predicts FPEPQ (dash-dotted line). Database size is varied by sampling; ×*n* indicates super-sampling by a factor *n* while */n* indicates sub-sampling by a factor *n*. This shows that Fold-seek *E*-values are substantially underestimated, diverging further from the ideal as database size increases. (b) Same plot for *E*-values corrected using Eq. 9, showing that this correction brings *E*-value estimates close to an ideal, comparable to Reseek (c), successfully accounting for variation in size. (c) Same plot for native Reseek *E*-values.

### 2.7 Dependence on alpha diversity

The abstract model shown in Fig. 3 gives the following expressions for *P*_Ø_(*s*), *P* (FP) and *P*_Ø_(*s*|FP) when there is no data-dependent filter,

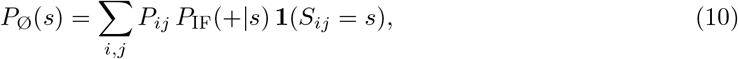

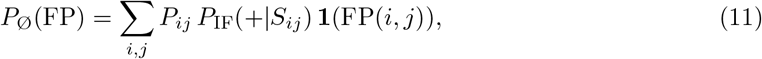

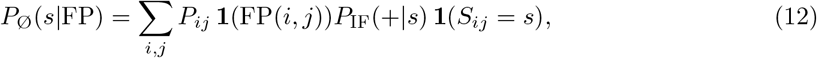

where *i* is a query structure, *j* is a target structure, *P*_*Q*_(*i*) is the probability of *i* in the query alpha diversity, *P*_*DB*_(*j*) is the probability of *j* in the database alpha diversity, *P*_*ij*_ = *P*_*Q*_(*i*)*P*_*DB*_(*j*) is the probability that the pair (*i, j*) will be sampled, *S*_*ij*_ is the score of aligning *i* to *j*, and **1**(*C*) is the so-called decision function which is one if the boolean condition *C* is true, otherwise zero. Then, by Bayes’ Theorem,

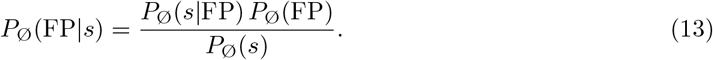

Eqs.(10 – 13) depend on alpha diversity via *P*_*ij*_, but do not depend on database size. We therefore expect no change in *P*_Ø_(FP|*s*) or *P*_Ø_(TP|*s*) = 1 − *P*_Ø_(FP|*s*) if we sub- or super-sample a database by domains, except for small fluctuations due to sampling effects. However, if we vary alpha diversity by sub-sampling superfamilies, we do expect to see a change. These predictions are confirmed by the results shown in Fig. 12.

### 2.8 Probability alignment is a TP

The Foldseek probability (prob column in outputs) is an estimate of *P* (TP|*s*) for a SCOP40 self-search. Supplementary Table S2 compares prob values reported by Foldseek to *P* (TP|prob) measured on a SCOP40 self-search. We find that prob is substantially overestimated on the training data; e.g. at prob= 0.999 the true probability is 0.730 and at prob= 0.900 the true probability is 0.444. With AFDB50 as the search database, our estimated E-value is *>* 10 for all Foldseek prob values ≤ 0.999.

### 2.9 Web calculator for log-linear E-values

We implemented a web page https://drive5.com/reseek/evalue_calculator.html which adds Ø_*LL*_ E-values to a user-supplied file containing TM scores, DALI Z scores or Foldseek E-values. A screenshot is shown in Fig. 11.

**Fig. 11:**
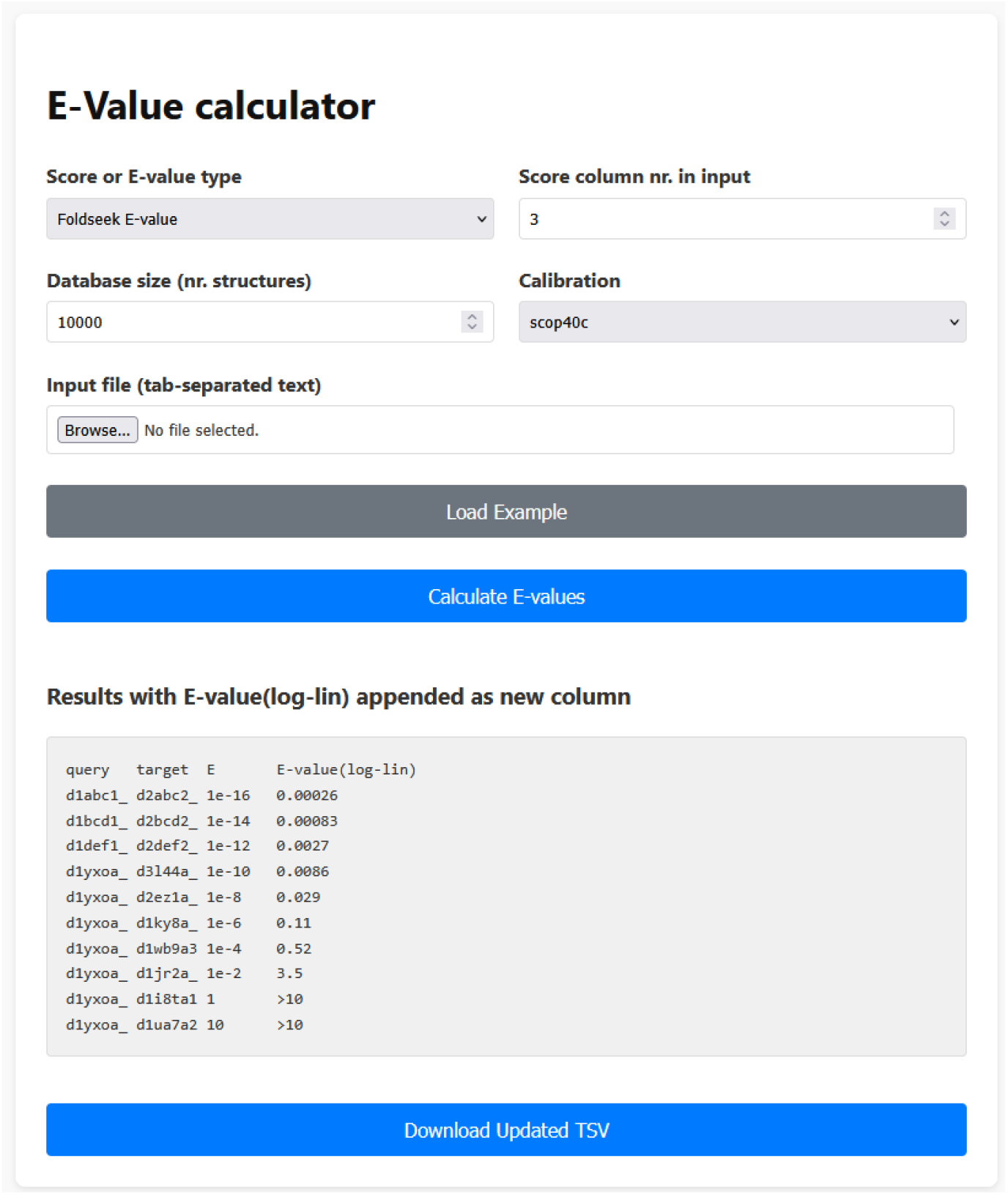
Web page for calculating E-values. Our calculator at https://drive5.com/reseek/evalue_calculator.html adds Ø_*LL*_ E-values to a user-supplied file containing TM scores, DALI Z scores or Foldseek E-values.

**Fig. 12:**
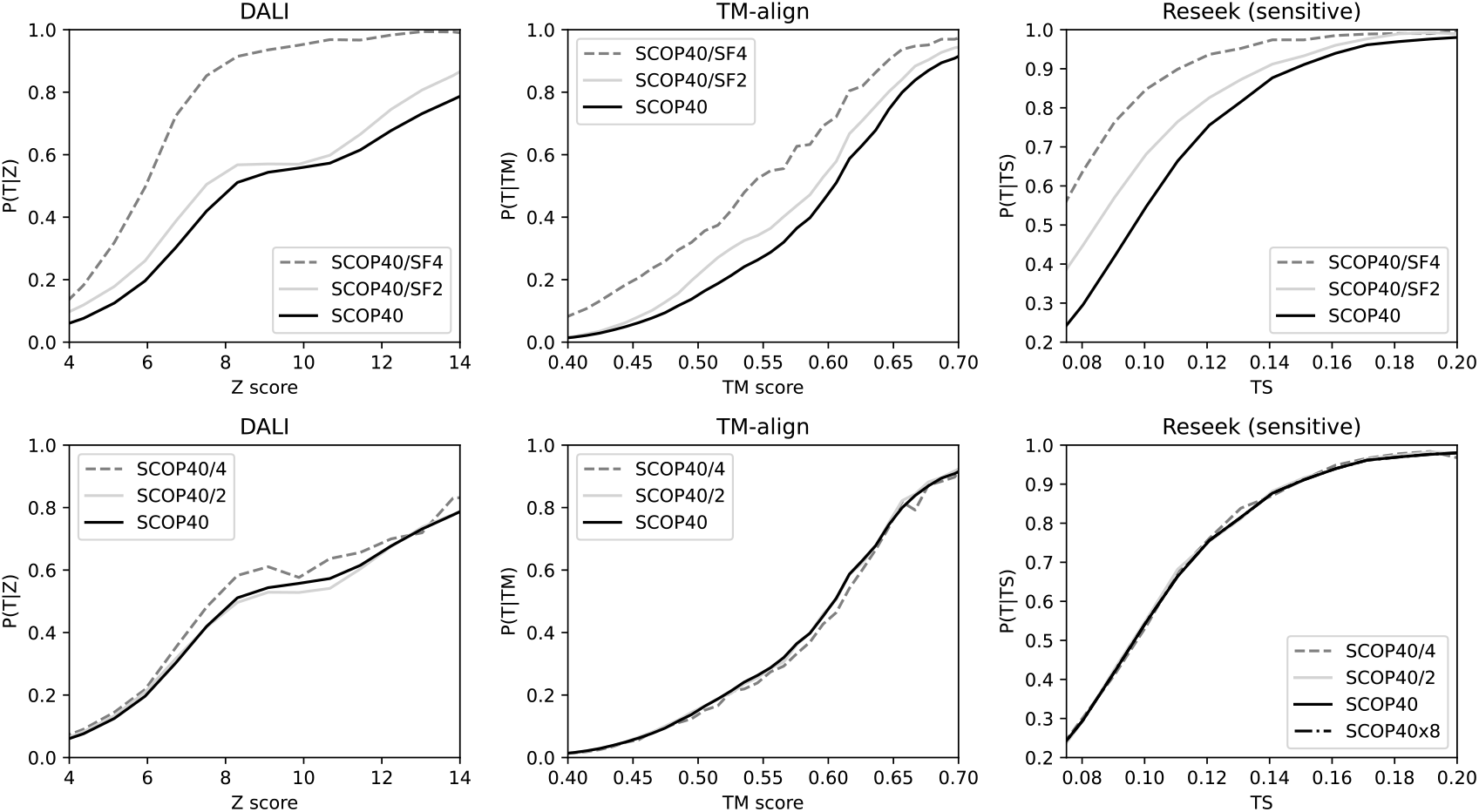
Distributions of *P* (TP|*s*) with varying references. Observed distributions of *P* (TP |*s*)—the ideal probability for assessing significance—for DALI Z-score (left), TM score (middle) and Reseek test statistic (TS, right) on all-vs-all alignments of SCOP40 Database size is varied by sampling; ×*n* indicates super-sampling by a factor *n, /n* indicates sub-sampling by a factor *n*, /SF*n* sub-sampling 1*/n* superfamilies, and ×*n* super-sampling by a factor *n* (×8 for Reseek only due to computational cost). The distributions vary strongly with alpha diversity (top row) but not with database size when alpha diversity is held fixed (bottom row). Thus, *P* (TP |*s*) = 1−*P* (FP |*s*) tends to vary with the search database, while *P* (*s* |FP) does not to a good approximation (Fig. 9).

## 3 Methods

### 3.1 Estimating E-values

Our goal is to calculate *E*(*s*), an estimate of the number of FP errors per query with score ≥ *s*. Let **D** be a database containing *D* structures; **Q** a query set with *Q* structures; *h*(*s*) the number of reported reported hits with score *s*; *H* = Σ _*s*_ *h*(*s*) the total number of reported hits, *P* (reported) = Σ _*s*_ *h*(*s*)*/Q* = *H/Q* the probability that a random query-target pair has a reported hit, or equivalently the number of reported hits divided by the number of possible hits; *P* (*s*) = *h*(*s*)*/H* the probability that a reported hit has score *s*; *P* (PF) the prior probability that a reported hit is a FP, *P* (*s*|FP) the probability that a false-positive reported hit has score *s*; and 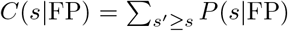 the probability that a false-positive hit has score ≥ *s*, i.e. the cumulative distribution of *P* (*s* | FP). We must consider that the reported hits will be a subset of all possible pairs if filters are used. By definition (estimation is implied as needed),

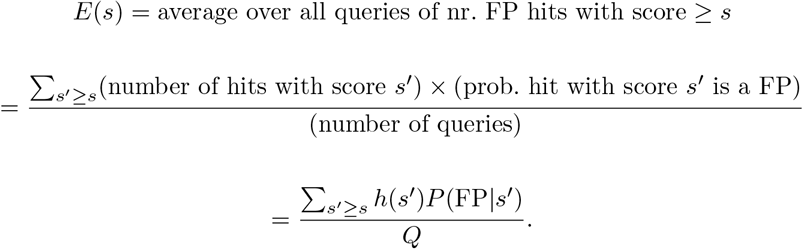

By Bayes’ Theorem,

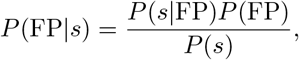

Giving

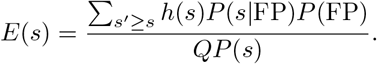

Substituting *P* (*s*) = *h*(*s*)*/H* then *h*(*s*) cancels and constants with respect to *s* can be moved outside the sum,

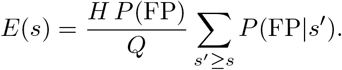

Noting that *H/Q* = *D P* (reported) and the sum is the cumulative distribution,

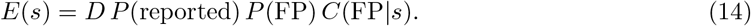

By comparison with Eq.(1) we see that the *p*-value is

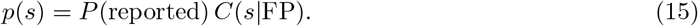

For a *comprehensive* algorithm such as TM-align, i.e. one which unconditionally reports an alignment for every query-target pair, *P* (reported) = 1 and we can assume *P* (FP) ≈ 1 because we expect that almost all query-target pairs are false positives in typical searches. Then Eq.(14) becomes

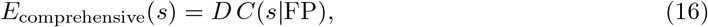

and the *p*-value is

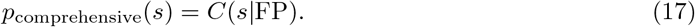

DALI discards alignments with *Z <* 2, which is a large majority of query-target pairs in a typical search. An alignment score threshold can be treated as an intrinsic filter because the score depends only on the pair being aligned (by contrast an *E*-value threshold is not intrinsic because *E* depends on the database size). Re-visiting Eq.(14), we should now keep in mind that all values are implicitly conditional on a hit being reported, i.e. having *Z* ≥ 2. Now *P* (reported) is generally ≪ 1; for example, in a SCOP40 all-vs-all search, DALI *P* (reported) = 0.037. A filter may discard a large fraction of false positives, which changes the prior expectation that a reported hit is a FP. When there is a filter, we propose that *P* (FP) = 1*/*2 is a reasonable default because the filter may discard a large majority of FPs and now we have no reason to suppose that FPs (or TPs) are more common in the hits. Then, for an algorithm with an intrinsic filter but no data-dependent filter, Eq.(14) becomes

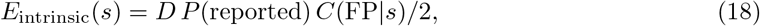

and the corresponding *p*-value is

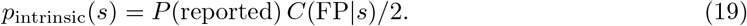

The value of *P* (reported) should ideally be measured on **D**, the search database used in practice. If the query set **Q** used in practice is large, then *P* (reported) can be measured on **Q** itself, otherwise by using a query dataset which is considered to be representative. Here we use SCOP40 by default, because it contains a diverse collection of structures, while noting that *P* (reported) is an unsupervised measurement and it is therefore not necessary to use a trusted reference. For example, if the query is a single viral capsid protein, then a large set of capsid proteins could be used to measure a query-specific *P* (reported).

### 3.2 Parameter fitting

Fitting to a potentially left-truncated Gumbel distribution was performed using the Pymoo package (Blank and Deb 2020). We implemented Ø_*LL*_ by log-linear fitting relevant distributions to the high-scoring tail using the Python `scipy.stats.linregress` function after reviewing a log-scaled plot to manually (i.e., visually) delineate a suitable range of scores in the tail. *E*-values reported by Ø_*LL*_ are pegged at *E* ≤ 10 because we found that distributions typically tend to deviate from the log-linear fit around *E* ~ 1.

#### 3.3 Software and databases

SCOP40 alignments for DALI and TM-align were obtained from supplementary data in (Van Kempen et al. 2024), which used SCOP40 v1.75; we therefore used this version of SCOP40 for other analyses here, including the construction of SCOP40c. For SCOP95 we used v2.08, the most recent version available at the time this work was done, to maximize the number of domains; we discarded the highly over-represented superfamily b.1.1 which contains antibodies, including many engineered variants. For CATH we used the Jan. 15th 2025 release clustered at 40% a.a. identity (CATH40), for PDB and we used the current Foldseek databases per Feb. 1st 2025, and for AFDB50 we used https://doi.org/10.5281/zenodo.14063905. Software versions were Foldseek v10-941cd33 and Reseek v2.6-e13c682.

## 4 Discussion

### 4.1 Protein structures are not random

Protein structures are not well-modeled by random sequences in the 3Di, *µ* or Mega alphabets. This reflects the fact that all alpha helices, all beta hairpins, all beta barrels…look alike, and similarity of these secondary and tertiary motifs is often due to convergence rather than homology (e.g. Fig. 7). For Foldseek and Reseek Smith-Waterman scores, the Gumbel distribution matches the bulk of the distribution, but fails to fit a “fat tail” of high-scoring FPs (Figs. 5, 6). We believe this fat tail is explained by convergent secondary and tertiary structure features, as illustrated by the alignment of unrelated domains shown in Fig. 7. This alignment has 39% 3Di sequence identity and score 992, placing it in the fat tail (Fig. 6, top) while a Gumbel fit predicts there should be none because *E*(992) = *D P*_Gumbel_(*s* ≥ 992|FP) ≪ 1, where *D* is the number of domains in SCOP40. Many of the 3Di letters for this pair of domains are in low-complexity regions according to segmasker (blue, lower-case); for example both domains end in `VV×××VVV` where × indicates a difference. In 3Di, V-rich regions are associated with alpha helices, which have locally similar conformations in unrelated proteins and may therefore be regarded as having convergent secondary structure. At a larger scale, the unmasked sequence (upper case, magenta) is comprised mainly of anti-parallel beta sheets, which also align well in their 3Di representation; note, for example, the motif `K×KIWTW××DPR×` (bold, shaded) where all columns contain identities or positive-scoring 3Di substitutions. Similarity in this larger-scale conformation (tertiary structure) is also convergent. While it may be possible to account for the fat tail by extending Karlin-Altschul statistics to incorporate low-complexity regions, we currently lack a suitable theoretical framework and therefore must resort to measuring distributions empirically. The high-scoring tail comprises a tiny fraction of the distribution, but this is the relevant fraction for distinguishing high-scoring false positives from true homologs in practice.

### 4.2 Gold standard reference

If theoretical distributions do not fit the data, then we need a trusted reference database to provide a representative sample of false positives for empirical fitting. Sequence homology can be inferred from protein structures, but there is no corresponding higher standard for structure—references such as CATH and SCOP necessarily infer homology using a combination of automated classification methods and human judgment, which is subject to some uncertainty; reasonable methods and reasonable algorithms may disagree. For our purposes in constructing a null model, an ideal reference should achieve a balance between false-positive and false-negative homology assignments, so that the measured distribution of FP scores according to the reference is close to correct despite the inevitable errors that will be present in the database. This goal conflicts with a conservative approach where structures are assigned to the same superfamily only when there is convincing evidence of homology. With SCOP, we attempted to mitigate these issues by creating SCOP40c, a curated subset of SCOP40 where, we believe, homology better corresponds to superfamilies (see Supplementary Material “Curating SCOP” and https://github.com/rcedgar/scop40c). In our judgment, SCOP40c is preferable to SCOP40 for training a null model and for benchmarking homology detection accuracy.

### 4.3 Universality of the FP distribution

We naively expect *P* (FP|*s*) to depend on alpha diversity per Eq.(13), potentially impeding modeling of large databases, which presumably have quite different diversities compared to a trusted reference database. Fortuitously, and perhaps surprisingly, we find that *P* (*s*|FP) is universal, i.e. independent of the reference to a good approximation (Fig. 9), enabling *P* (FP|*s*) to be calculated via Bayes’ Theorem. We believe this can be explained by a universal tendency for proteins to form similar secondary and tertiary structures.

### 4.4 Significance of TM score and the TM=0.5 cutoff

Previous work (Xu and Zhang 2010) showed that the bulk of the TM score FP distribution is approximately Gumbel, then used the fitted Gumbel to estimate *p*-values, reporting e.g. that *p* = 5.5 × 10^*−*7^ for TM=0.5. Revisiting this result, we find that 167,216 / 124,638,772 FP pairs in SCOP40 have TM *>* 0.5, giving a measured *p*-value of 1.3 × 10^*−*3^ using fold as a truth standard without fitting or approximation. Thus, the *p*-value for TM=0.5 given by (Xu and Zhang 2010) is underestimated by more than three orders of magnitude due to the poor fit of the Gumbel distribution in the high-scoring tail (our Fig. 5). The analysis in (Xu and Zhang 2010) concludes that “…it seems quite safe to assign TM-score = 0.5 as a rough but quantitative cutoff for protein Fold/Topology definition, i.e. most of proteins with TM-score *>* 0.5 can be considered as of the same topology whereas most proteins with a TM-score *<* 0.5 should be of different topology”. We disagree with this conclusion. In our opinion, fixed score thresholds are generally unsafe and should be replaced by database-specific *E*-value thresholds. For this case, we find *P* (same fold|TM ≥ 0.5) = 0.75 (note again this is without approximation or fitting), so the guideline “TM ≥ 0.5 suggests same fold” has a 25% chance of failing in a single test on SCOP40. For testing many alignments, as in a database search, this would surely allow many FPs, as reflected by the Ø_*LL*_ estimate that the TM=0.5 has *E >* 10 on any database comparable in size to SCOP40 or larger (Supp. Table S1). Also, more than half (531,939 out of 1,036,538) of pairs with the same fold have TM *<* 0.5, so the guideline “TM *<* 0.5 suggests different fold” fails to recognize more than 50% of true positives.

### 4.5 Practical importance of accurate E-values

The TP and FP distributions for TM score on SCOP40 are shown in Fig. 1. Notice that the bulk of the TP distribution overlaps the FP distribution, and the peak of the FP distribution is orders of magnitude larger than the TP distribution. This illustrates the general point that a score cutoff cannot cleanly separate TPs and FPs—it is necessary to make a choice between high precision (reducing FPs) or high sensitivity (reducing FNs) because these are strongly conflicting goals. The gold standard for making this trade-off is to set an *E*-value threshold to limit the expected number of FPs. Sensitivity and precision at a given score cutoff depend on diversity, and the expected number of errors increases with database size. A null model which accurately accounts for diversity, database size and convergent evolution of structural features is therefore essential for robust estimation of an *E*-value.

### 4.6 Fold as a truth standard

Folds are arbitrary in the sense that it is not possible in principle to determine whether two domains have the same fold without referring to a particular classification scheme. From this perspective, folds are similar to taxonomic clades at a given rank, which may be lumped or split according to taste, except that clades should be homologous, while with folds, there is no analogous constraint. By contrast to folds, superfamilies are not arbitrary because homology is objectively defined. The universality of the FP distribution that we observe in practice suggests that superfamilies in SCOP and CATH are reasonably close to correct, and also that it is reasonable to consider a generic alignment score as a basis for binary classification with superfamily as a standard. Evolution, and hence homology detection, can be universally modeled across all proteins using a single set of parameters (e.g. by a BLOSUM matrix (Henikoff and Henikoff 1992)), while folds and topologies as currently defined cannot be universally modeled. Given these considerations, it seems clear that different cutoffs should be applied to different categories of fold, and such cutoffs will be at best very rough discriminators. Robust fold discrimination, if indeed this is possible, will necessarily require a multiclass classifier trained on a particular scheme such as SCOP or CATH. A binary classifier that attempts to distinguish same from different folds via a single generic score threshold, such as TM=0.5, cannot be robust.

### 4.7 Significance of DALI Z score

DALI reports a Z score, which is a standard measure of significance in statistics, as opposed to other scores such as Smith-Waterman or TM-score, which are not significance measures. Note that in this work, we use “significance” exclusively to mean a quantitative estimate of error probability or number of errors. Z is calculated from a raw score obtained by aligning contact maps (see (Holm and Sander 1993), Eq.(1)). This raw score is used internally by DALI but otherwise rarely mentioned in practice. Z assumes a Gaussian distribution and is expressed as a number of standard deviations of the raw score from the mean (Moore 2010). For example, Z=5 implies “five-sigma significance”, i.e. the alignment score is five standard deviations higher than the mean, corresponding to a *p*-value of 2.9 ×10^*−*7^ (Lyons 2013). However, we find that Z=5 corresponds to a *p*-value of 0.1 using SCOP40 calibration or 0.39 using SCOP40c (Supp. Table S1), showing that DALI’s reported *Z* ~ 5 overestimates significance by roughly six orders of magnitude.

### 4.8 Foldseek E-values

We found that Foldseek also substantially overestimates significance. For example, we find that a reported *E*-value of 10^*−*6^ corresponds to a measured FPEPQ of ~ 10^*−*1^ on SCOP40 (Supp. Table 1), an underestimate of five orders of magnitude. This was measured directly on SCOP40, the same reference used by Foldseek for calibration. Extrapolating to larger databases (Eq. 9, Supp. Table 1), we project that a Foldseek *E*-value of 10^*−*6^ corresponds to FPEPQ ~ 2 on both PDB (~ 10^6^ structures) and AFDB50 (~ 50× 10^6^ structures), i.e. roughly six orders of magnitude underestimated. The underestimate on SCOP40 is largely explained by the assumption of a Gumbel distribution, which does not fit raw scores well, especially in the high-scoring tail (Fig. 8). If sequences were random, then the Smith-Waterman scores for the query (*SW*_*q*_) and for the reversed query (*SW*_*qrev*_) would each be well-modeled by a Gumbel distribution. However, the raw score is the difference between these scores, *s*_*raw*_ = *SW*_*q*_ − *SW*_qrev_. The difference between two random variables sampled from a Gumbel has a logistic PDF (McFadden 1972), and under a random sequence model *s*_*raw*_ should therefore be fit to logistic rather than to Gumbel. Thus, on SCOP40 the underestimate is primarily explained by fitting to an inappropriate distribution, and with larger databases the scaling applied by Foldseek (Eq.(4)) does not extrapolate accurately, as shown by the increasingly underestimated significance with database size (Fig. 10).

## 5 Conclusion

We have described and validated a new approach to quantifying statistical significance for protein structure searches, enabling principled estimates of *p*-values and *E*-values. It is conceptually more complicated than previous approaches, necessitated by the essentially non-random nature of protein structures and the requirement to model recent algorithm innovations (filters), yet enables the calculation of an *E*-value from an alignment score in a simple expression requiring only two fitted parameters, or four if there is a filtering step. We show that this approach successfully accounts for variations in database size and diversity, and can therefore be applied with confidence to the burgeoning protein structure databases unleashed by the AI folding era.

## Data and code availability

Code and data are deposited at https://github.com/rcedgar/null_model2 and https://github.com/rcedgar/scop40c under the GPL3 license.

