## Supplementary Material for "Protein structure alignment significance is often exaggerated" (from another version of this preprint: version 1, 2025-07-19)

### 1 Curated SCOP40

We identified issues with SCOP40 which motivated us to construct a curated subset. There are many short folds with simple conformations where we saw no reasonable basis for deciding whether similarity is due to homology or convergence (Fig. S6). We found several distinct folds which are clearly homologous, having similar conformations and unambiguous sequence similarity (examples in Figs. S7 and S8). We found other cases where three or four of our chosen methods (DALI, TM-align, Foldseek and Reseek) agreed with high confidence that different superfamilies are homologous. We developed an automated pipeline to delete folds with these issues, giving a curated subset (SCOP40c) with 8,340/11,211 domains, 1,438/1,960 superfamilies and 1,005/1,194 folds from SCOP40. Code and data deposited at `https://github.com/rcedgar/scop40c`.

### 2 Truth standards

Six distinct SCOP-based truth standards from the literature, denoted Fam1 through Fold6, are described in the supplementary material of (Edgar, 2024). In this work, we use SF2, which considers an alignment as true positive (TP) if the structures belong to the same superfamily (SF), otherwise false positive (FP). This differs from SF3, the standard (Brenner et al., 1998) which is most commonly used, because SF3 ignores alignments between different superfamilies in the same fold (DSSF) for which is considered ambiguous. On SCOP40, there are 581,772 DSSF pairs, more than the number of TPs (454,766). As seen in Fig. S5, in the high-scoring tail which is most relevant to statistical significance, DSSFs comprise the majority of pairs (gray). We agree that homology is uncertain for DSSF pairs, but we believe that they are mostly not homologous or are so far diverged in function that they are effectively FPs for most searches. Therefore, we prefer SF2 because we believe that the common practice of ignoring

DSSF pairs substantially underestimates error rates.

### 3 Foldseek patch to report Smith-Waterman and raw score

We modified the Foldseek source code to report Smith-Waterman and raw scores by editing `src/strucclustutils/structurealign.cpp` in `https://github.com/steineggerlab/foldseek` (commit `941cd33`). The patched version of this source file is found at `foldseek_src_patch/structurealign.cpp` in `https://github.com/rcedgar/null_model2`.

### Supplementary Tables and Figures

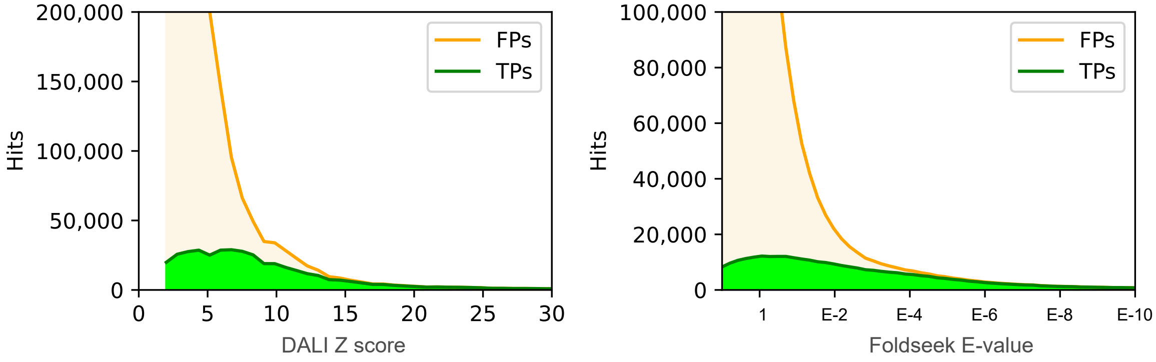


**Figure S1.** TP and FP distributions for DALI and Foldseek
See also Fig. 1 in the main text.

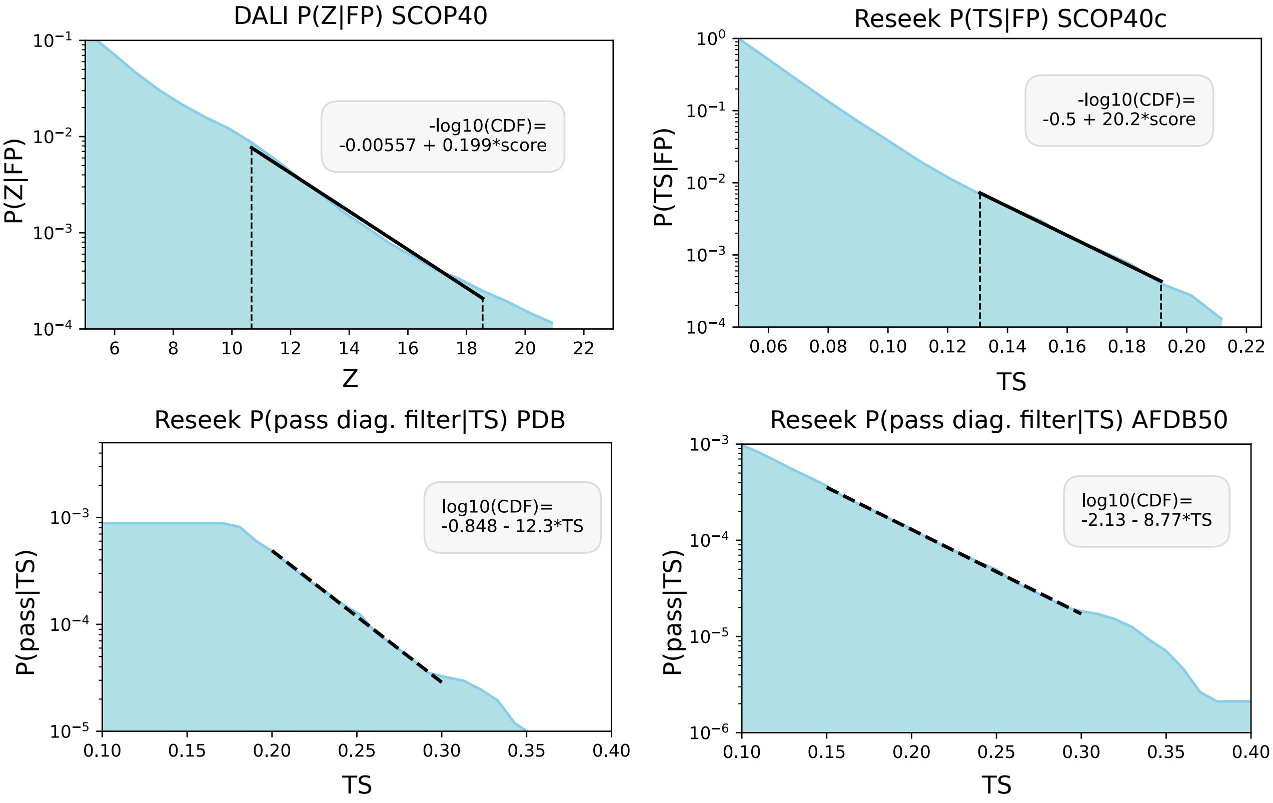


**Figure S2.** Representative log-linear fits
We found empirically that relevant distributions were well approximated by log-linear regression in the high-scoring regime of interest here, as illustrated by the examples shown in this figure.

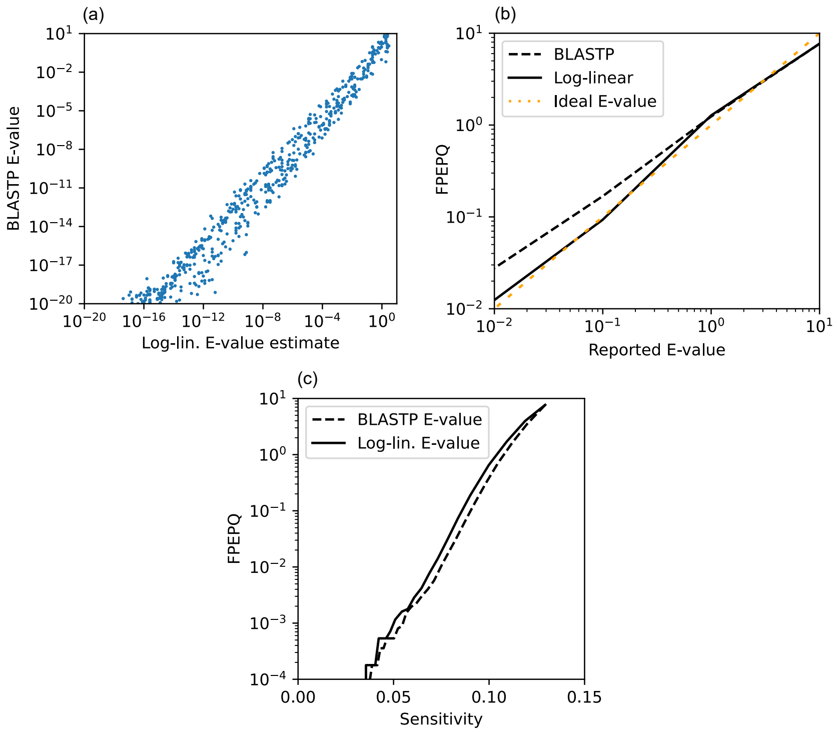


**Figure S3.** Validation on BLASTP
We applied Ø$_{LL}$ to BLASTP raw scores, with results for SCOP40 shown in this figure. (a) Scatterplot of a random sample of 1,000 $E$-values with Ø$_{LL}$ estimate on the $x$ axis and native BLASTP $E$-value on the $y$ axis. (b) $E$-value (predicted FPEPQ) on the $x$ axis against measured FPEPQ on the $y$ axis. Both the native BLASTP $E$-value and the Ø$_{LL}$ $E$-value are close to ideal, with Ø$_{LL}$ closer to the ideal. (c) Sensitivity vs. FPEPQ plot showing similar accuracy for Ø$_{LL}$ and native $E$-values.

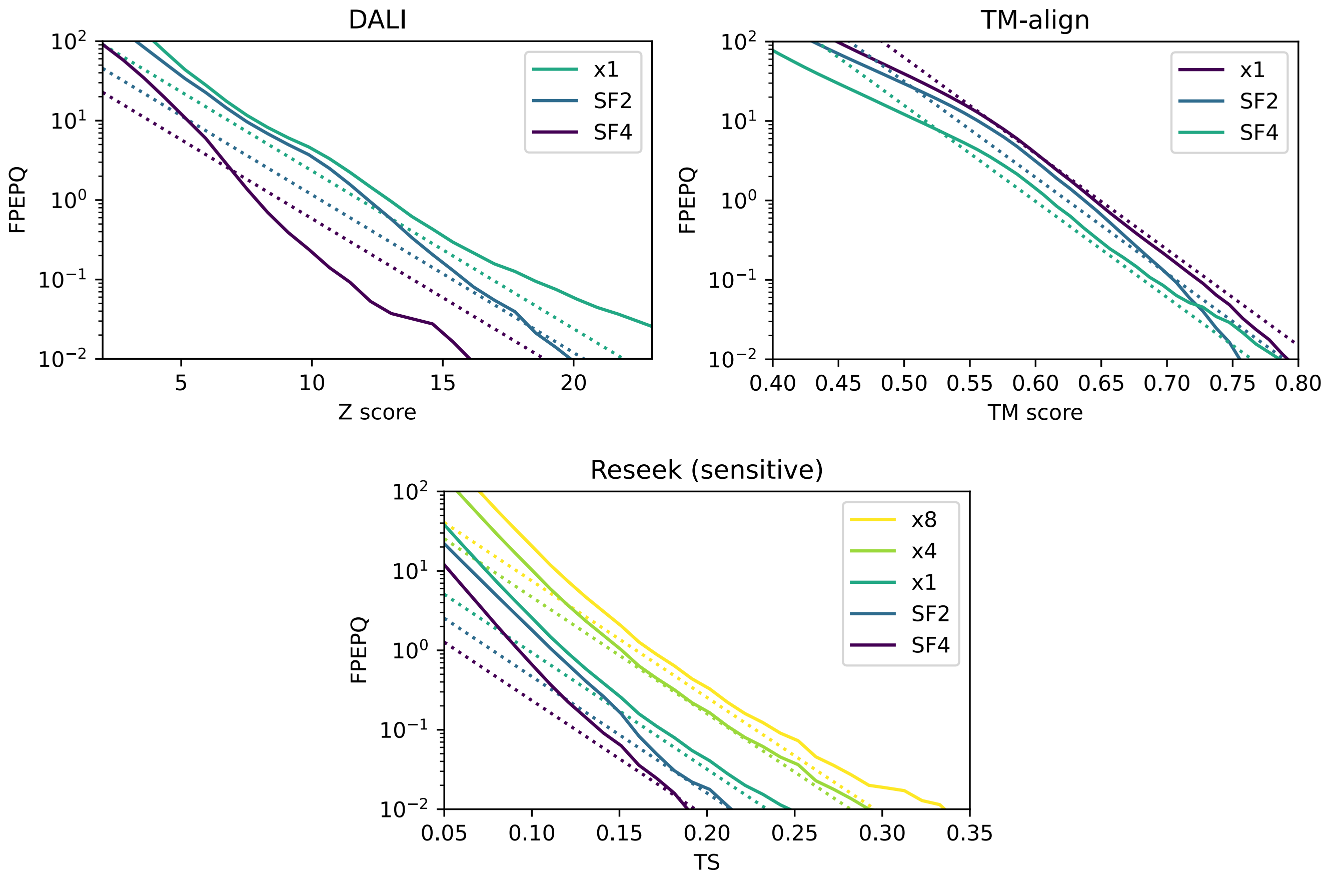


**Figure S4.** Validation on DALI, TM-align and Reseek
We applied $\text{Ø}_{LL}$ to DALI Z-scores, TM scores and the Reseek test statistic (TS) with results shown in this figure. Solid lines are measured FPEPQ as a function of score, dotted lines are $\text{Ø}_{LL}$ $E$-values (estimates of FPEPQ). Results are shown for sub-sampling by superfamily (SF2 and SF4) and super-sampling by factors 4 and 8 ($\times 4$ and $\times 8$, respectively; for Reseek only due to the high cost of DALI and TM-align). These results show that $\text{Ø}_{LL}$ accounts well for systematic differences in FPEPQ induced by differences in alpha diversity and database size.

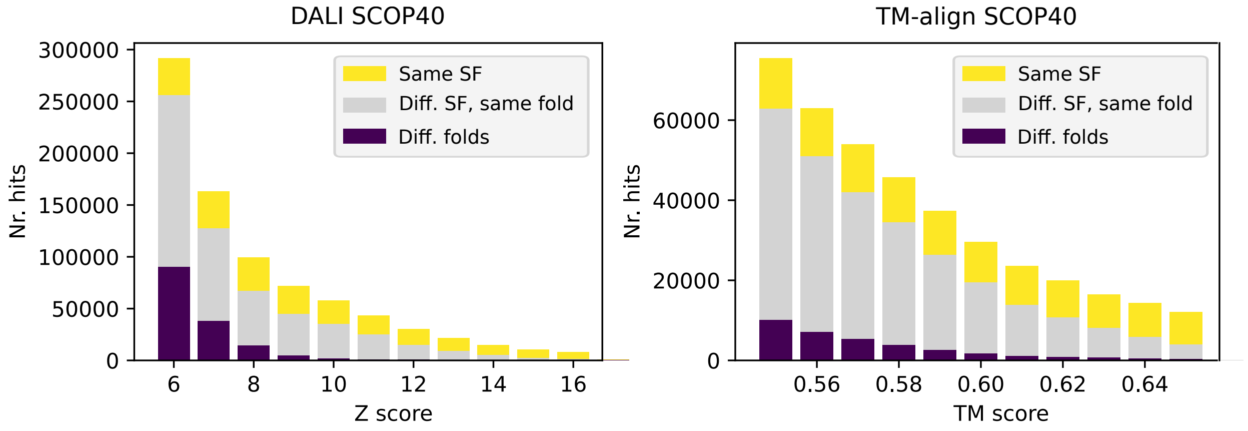


**Figure S5.** Comparison of truth standards
Stacked histograms show the numbers of alignments in the same superfamily (yellow), different superfamily in the same fold (DSSF, gray), and different folds (purple), respectively, using SCOP40 as the reference, binned by DALI Z-score (left) and TM score (right). The range of scores shown is the high-scoring tail where discriminating TPs from FPs is most challenging. Note that this range is dominated by DSSF alignments which are ignored by the truth standard which is commonly used in the literature. We believe that most DSSF alignments are not homologous or are so diverged in function that they are effectively FPs for most searches. We therefore consider alignments to be TPs if domains are in the same superfamily, otherwise FPs.

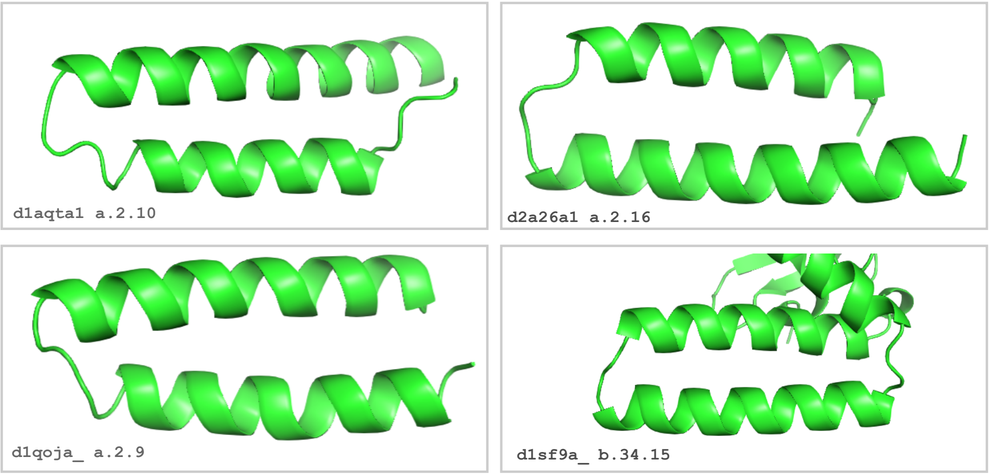


**Figure S6.** Short domains classified to different superfamilies by SCOP
This figure shows cartoon representations of domains d2a26a1, d1aqta1 and d1qoja_ which have similar alpha-hairpin conformations and are classified to superfamilies a.2.16, a.2.10 and a.2.9, respectively, by SCOP. In our opinion, the conformations of these domains are sufficiently similar that convergence and homology are both plausible explanations for the observed similarity. As illustrated by domain d1sf9a_ (bottom-right corner), which is classified to fold b.34 (assigned to class b, which nominally contains all-beta structures), alpha hairpins are common motifs. In SCOP40c, we deleted all domains $\leq$ 80 a.a. and all members of superfamilies with median $\leq$ 80 a.a. to reduce the number of cases where homology is ambiguous due to short length.

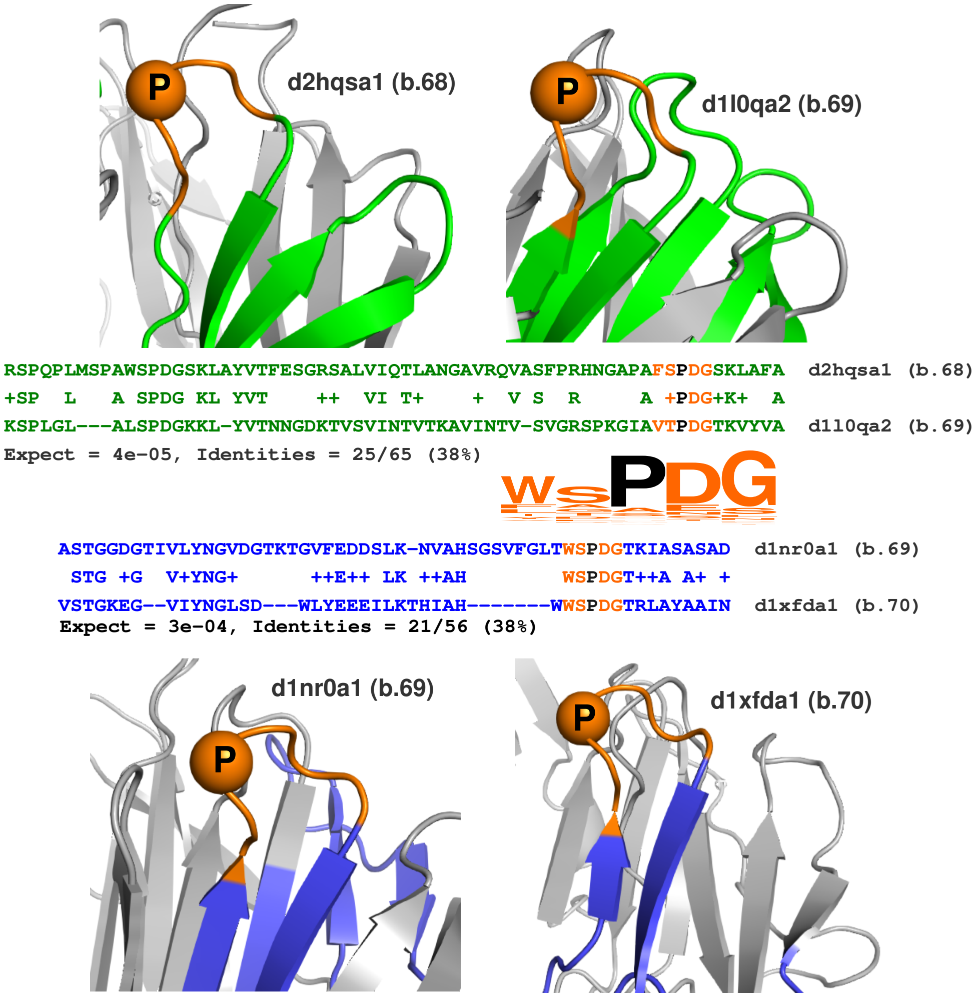


**Figure S7.** Homologous domains classified to different folds by SCOP
This figure shows domains which have clear similarity in both structure and sequence, but are classified to three different folds by SCOP. All three folds are beta-propellers, distinguished by the number of blades in the propeller: six in fold `b.68`, seven in `b.69` and eight in `b.68`. Two pair-wise BLASTP alignments are shown, both of which have 38% sequence identity between different folds and highly significant $E$-values $4 \times 10^{-5}$ and $3 \times 10^{-4}$, respectively ($E$-values calculated assuming a database of SCOP40 size). The sequence motif `WSPDG` is well-conserved across all folds; the sequence logo shown here was obtained from a Muscle-3D (Edgar and Tolstoy, 2024) multiple structure alignment for domains in all three folds. A conserved `P` is exposed at the turn in a beta hairpin which connects two blades in the propeller. We removed these folds, and other folds having high-confidence homology, from our curated reference (SCOP40c).

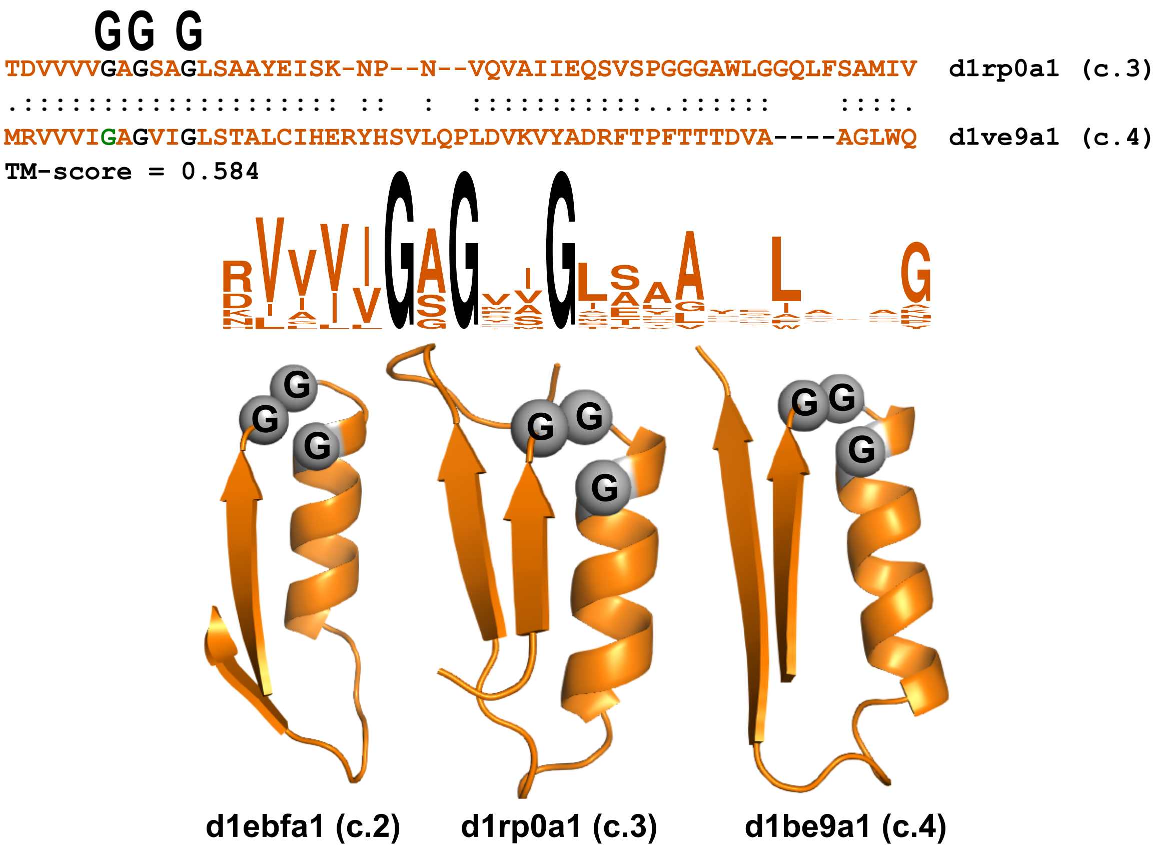


**Figure S8.** Homologous domains classified to different folds by SCOP
Three domains which have clear similarity in both structure and sequence, but are classified to three different folds by SCOP: `c.1`, `c.2` and `c.3`, respectively. Top is an cross-fold alignment by TM-align showing a region where most aligned residues are < 5Å (shown by ':'). Middle is a sequence logo derived from a Muscle-3D alignment of collected domains in these folds, showing well-conserved positions with three perfectly conserved `G`s. Bottom are cartoon representations showing a conserved conformation with an alpha helix parallel to a beta hairpin. We removed these folds, and other folds having high-confidence homology, from our curated reference (SCOP40c).

| DALI Z-score | P-val | P-valc | SCOP40 | SCOP40c | PDB | PDBc | AFDB50 | AFDB50c |
|---|---|---|---|---|---|---|---|---|
| 5 | 1.0e-01 | 3.9e-01 | >10 | >10 | >10 | >10 | >10 | >10 |
| 10 | 1.0e-02 | 6.8e-03 | 2.3e+00 | 1.5e+00 | >10 | >10 | >10 | >10 |
| 15 | 1.1e-03 | 1.2e-04 | 2.4e-01 | 2.7e-02 | >10 | 2.1e+00 | >10 | >10 |
| 20 | 1.1e-04 | 2.1e-06 | 2.4e-02 | 4.7e-04 | 1.9e+00 | 3.7e-02 | >10 | 2.3e+00 |
| 25 | 1.1e-05 | 3.7e-08 | 2.4e-03 | 8.3e-06 | 1.9e-01 | 6.6e-04 | >10 | 4.0e-02 |
| 30 | 1.1e-06 | 6.5e-10 | 2.5e-04 | 1.5e-07 | 1.9e-02 | 1.2e-05 | 1.2e+00 | 7.0e-04 |
| 35 | 1.1e-07 | 1.1e-11 | 2.5e-05 | 2.6e-09 | 2.0e-03 | 2.0e-07 | 1.2e-01 | 1.2e-05 |
| 40 | 1.1e-08 | 2.0e-13 | 2.5e-06 | 4.5e-11 | 2.0e-04 | 3.6e-09 | 1.2e-02 | 2.2e-07 |
| 45 | 1.1e-09 | 3.5e-15 | 2.6e-07 | 8.0e-13 | 2.0e-05 | 6.3e-11 | 1.2e-03 | 3.8e-09 |
| 50 | 1.2e-10 | 6.2e-17 | 2.6e-08 | 1.4e-14 | 2.1e-06 | 1.1e-12 | 1.2e-04 | 6.7e-11 |
| 55 | 1.2e-11 | 1.1e-18 | 2.6e-09 | 2.5e-16 | 2.1e-07 | 1.9e-14 | 1.3e-05 | 1.2e-12 |
| 60 | 1.2e-12 | 1.9e-20 | 2.7e-10 | 4.3e-18 | 2.1e-08 | 3.4e-16 | 1.3e-06 | 2.1e-14 |

| TM-align TM score | P-val | P-valc | SCOP40 | SCOP40c | PDB | PDBc | AFDB50 | AFDB50c |
|---|---|---|---|---|---|---|---|---|
| 0.50 | 5.6e-03 | 6.4e-03 | >10 | >10 | >10 | >10 | >10 | >10 |
| 0.55 | 1.4e-03 | 1.3e-03 | >10 | >10 | >10 | >10 | >10 | >10 |
| 0.60 | 3.5e-04 | 2.7e-04 | 3.9e+00 | 3.0e+00 | >10 | >10 | >10 | >10 |
| 0.65 | 8.6e-05 | 5.4e-05 | 9.7e-01 | 6.1e-01 | >10 | >10 | >10 | >10 |
| 0.70 | 2.2e-05 | 1.1e-05 | 2.4e-01 | 1.2e-01 | >10 | 9.8e+00 | >10 | >10 |
| 0.75 | 5.4e-06 | 2.3e-06 | 6.0e-02 | 2.5e-02 | 4.7e+00 | 2.0e+00 | >10 | >10 |
| 0.80 | 1.3e-06 | 4.6e-07 | 1.5e-02 | 5.2e-03 | 1.2e+00 | 4.1e-01 | >10 | >10 |
| 0.85 | 3.3e-07 | 9.4e-08 | 3.7e-03 | 1.1e-03 | 2.9e-01 | 8.3e-02 | >10 | 5.0e+00 |
| 0.90 | 8.3e-08 | 1.9e-08 | 9.3e-04 | 2.2e-04 | 7.3e-02 | 1.7e-02 | 4.4e+00 | 1.0e+00 |
| 0.95 | 2.1e-08 | 3.9e-09 | 2.3e-04 | 4.4e-05 | 1.8e-02 | 3.5e-03 | 1.1e+00 | 2.1e-01 |

| Foldseek E-value | P-val | P-valc | SCOP40 | SCOP40c | PDB | PDBc | AFDB50 | AFDB50c |
|---|---|---|---|---|---|---|---|---|
| 1.0e+01 | 1.2e-01 | 5.7e-01 | >10 | >10 | >10 | >10 | >10 | >10 |
| 1.0e+00 | 4.8e-02 | 1.2e-01 | >10 | >10 | >10 | >10 | >10 | >10 |
| 1.0e-01 | 1.9e-02 | 2.5e-02 | >10 | >10 | >10 | >10 | >10 | >10 |
| 1.0e-02 | 7.6e-03 | 5.3e-03 | 3.7e+00 | 2.6e+00 | >10 | >10 | >10 | >10 |
| 1.0e-03 | 3.0e-03 | 1.1e-03 | 1.4e+00 | 5.2e-01 | >10 | >10 | >10 | >10 |
| 1.0e-04 | 1.2e-03 | 2.4e-04 | 5.7e-01 | 1.1e-01 | >10 | 2.2e+00 | >10 | >10 |
| 1.0e-05 | 4.8e-04 | 5.0e-05 | 2.5e-01 | 2.6e-02 | 4.9e+00 | 5.1e-01 | >10 | 8.3e+00 |
| 1.0e-06 | 1.9e-04 | 1.1e-05 | 1.2e-01 | 6.6e-03 | 2.3e+00 | 1.3e-01 | >10 | 2.1e+00 |
| 1.0e-07 | 7.6e-05 | 2.2e-06 | 5.9e-02 | 1.8e-03 | 1.2e+00 | 3.4e-02 | >10 | 5.6e-01 |
| 1.0e-08 | 3.0e-05 | 4.7e-07 | 3.1e-02 | 4.9e-04 | 6.1e-01 | 9.5e-03 | 9.9e+00 | 1.6e-01 |
| 1.0e-09 | 1.2e-05 | 1.0e-07 | 1.7e-02 | 1.4e-04 | 3.3e-01 | 2.7e-03 | 5.4e+00 | 4.4e-02 |
| 1.0e-10 | 4.8e-06 | 2.1e-08 | 9.3e-03 | 4.1e-05 | 1.8e-01 | 8.0e-04 | 2.9e+00 | 1.3e-02 |

**Table S1** P-values and $E$-values by log-linear fits. This table shows $p$-values and $E$-values estimated by $\emptyset_{LL}$ for DALI, TM-align and Foldseek. Pval is $p$-value with SCOP40 as reference, Pvalc with SCOP40c as reference. $E$-values according to $\emptyset_{LL}$ are given for search databases SCOP40, PDB and AFDB50; c suffix indicates that SCOP40c was used as reference, SCOP40 otherwise.

| prob | P(TP\|prob) | $E_{\text{foldseek}}$ | FPEPQ | $\text{Ø}_{LL}$ SCOP40 | $\text{Ø}_{LL}$ AFDB50 |
|---|---|---|---|---|---|
| 0.999 | 0.730 | 0.0012 | 0.17 | 0.18 | >10 |
| 0.949 | 0.495 | 0.01 | 0.35 | 0.43 | >10 |
| 0.900 | 0.444 | 0.015 | 0.71 | 0.5 | >10 |

**Table S2** Significance of high Foldseek prob values.
The table shows `prob` values 0.999, 0.949 (the closest reported value to 0.95) and 0.900. The `prob` field is described as the probability that a hit in a SCOP40 all-vs-all search is a TP. $P(\text{TP}|\text{prob})$ is the measured value for this probability as the fraction of SCOP40 alignments with this value of prob which are TPs. $E_{\text{foldseek}}$ is the median E-value reported by Foldseek for hits with the given prob value. FPEPQ is the measured false positive errors per query with prob $\geq$ the given value. $\text{Ø}_{LL}$ SCOP40 is the E-value estimated by ØLL for SCOP40 as the search database. $\text{Ø}_{LL}$ AFDB50 is is the E-value estimated by $\text{Ø}_{LL}$ for AFDB50 as the search database, which is $> 10$ for all prob values $\leq 0.999$.
